# Angulin-1/LSR inhibition transiently disrupts the blood-tumor barrier to enhance doxil permeability and impair malignant glioma progression

**DOI:** 10.1101/2025.07.31.667901

**Authors:** Dominique Ferguson, Minhye Kwak, Sanghee Lim, Melissa Cesaire, Jatia Mills, Mahalia Dalmage, Jane Jones, Sergey Tarasov, Marzena Dyba, Rob Robey, Yanbo Yang, Shae K. Simpson, Baktiar Karim, Donna Butcher, Robyn Gartrell, Michael Gottesman, Sadhana Jackson

## Abstract

The blood-tumor barrier (BTB) prevents effective central nervous system (CNS) drug delivery, especially in malignant gliomas. Brain endothelium predominates the BTB and connects through bicellular and tricellular tight junctions (TJ). Angulin-1/LSR, is a highly expressed endothelial tricellular TJ. Our studies explore the role of Angubindin-1, an Angulin-1/LSR binder, to disrupt tricellular TJ integrity, increase drug entry and hamper glioma progression. Using rat brain endothelial cells (RBMVEC) we tracked Angulin-1/LSR localization and expression to the membrane; binding tightest to Angubindin-1 2-8 hours post-treatment (*p* < 0.05). Angubindin-1 dose-dependently reduced bicellular and tricellular TJs 1-4 hours post treatment (*p* < 0.05), returning to baseline by 24 hours (*p* < 0.05). In human and rat-derived glioma cells, Angubindin-1 transiently reduced Angulin-1/LSR expression between 2-8 hours (*p* < 0.05), with return to baseline by 24 hours (*p* < 0.001). Silenced Angulin-1/LSR expression on endothelium resulted in decreased mRNA levels of bicellular (occludin, claudin-5, ZO-1) and tricellular (tricellulin/MARVELD2, angulin-1/LSR) TJs compared to control (*p* < 0.01). Angubindin-1 treatment also inhibited efflux transporter P-gp in both RBMVECs and glioma cells with high P-gp expression only. Orthotopic rat glioma models were treated with Doxil (3 mg/kg), Angubindin-1 (10 mg/kg), or combination to evaluate BTB permeability/drug accumulation, and overall survival. Combination therapy enhanced Doxil tumor accumulation by 20% (*p* < 0.001), reduced tumor volume by day 14 (77.5% vs. 81.6%, *p* < 0.05), and significantly extended survival compared to Doxil alone (24 days vs. 18 days, *p* < 0.0001). These findings demonstrate the effects of tricellular tight junction inhibition on disrupting the BTB, enhancing CNS drug delivery, and improving rodent glioma survival.

**Significance:** This study demonstrates that Angubindin-1, a targeted modulator of tricellular tight junction protein Angulin-1/LSR, transiently disrupts BTB integrity to enhance chemotherapy delivery and prolong survival in glioma-bearing rats.

**Graphical Abstract:** 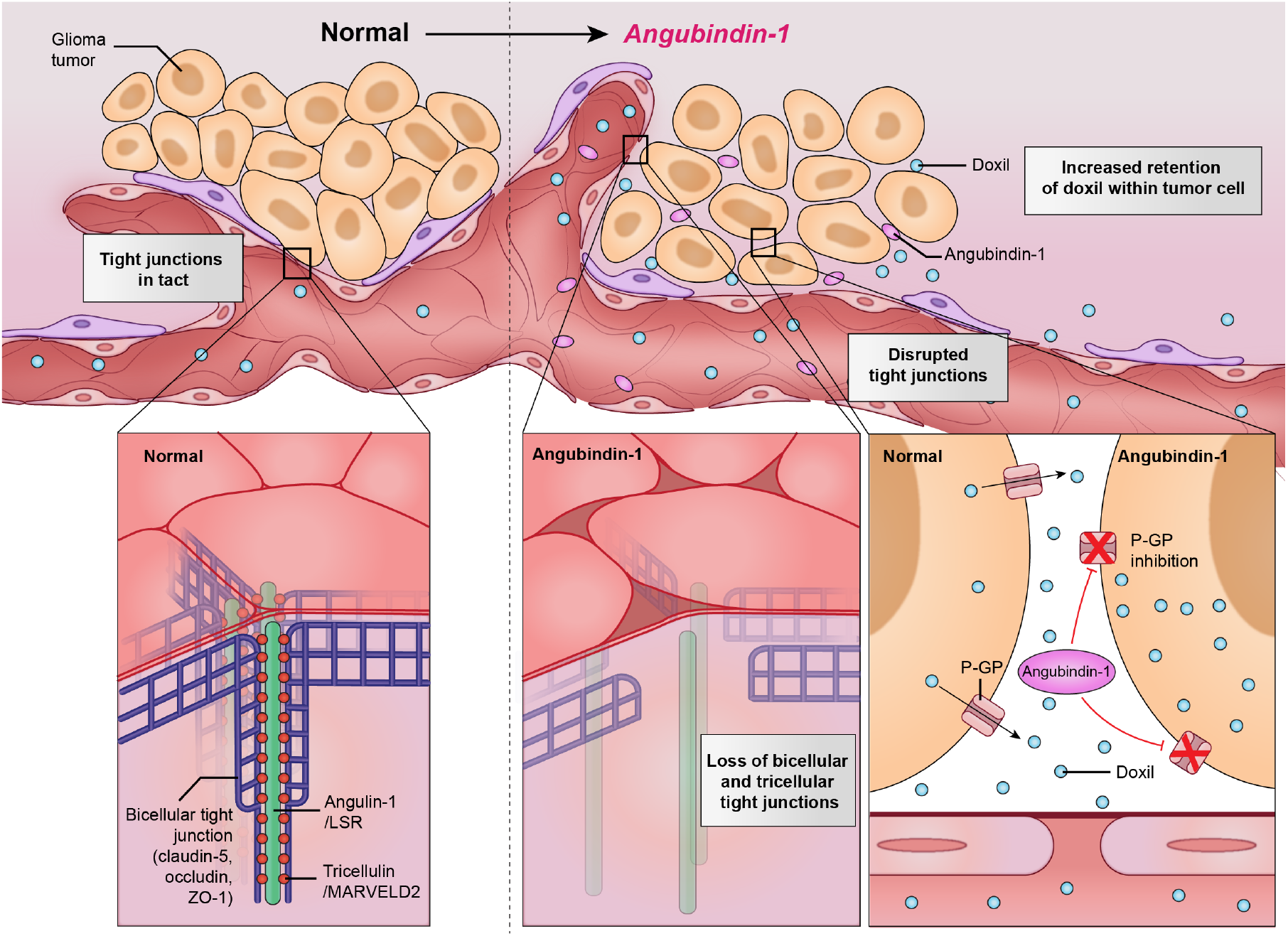

Angubindin-1 targets both bicellular tight junctions and the tricellular tight junction protein, Angulin-1/LSR, in brain endothelial and glioma cells leading to transient disruption of the blood–tumor barrier (BTB) and inhibition of P-glycoprotein towards enhanced Doxil penetration and reduced tumor burden.

## Introduction

The brain endothelium plays a critical role in regulating drug delivery to the central nervous system (CNS), particularly in the context of malignant tumors such as Isocitrate Dehydrogenase (IDH) wild-type (WT) glioblastoma (GBM). In these tumors, the endothelial vasculature presents a significant barrier to therapeutic delivery, both within and beyond the tumor core [1-6]. The integrity of the brain endothelium is primarily maintained by tight junction (TJ) complexes composed of both bicellular (e.g., claudin-5, zonula occludens-1, occludin) and tricellular TJ proteins (e.g., angulin-1 and tricellulin/MARVELD2) [7, 8]. These junctions, along with specialized efflux transporters such as P-glycoprotein (P-gp), work in concert to regulate paracellular and transcellular permeability; limiting access of xenobiotics and therapeutics to the brain.

P-glycoprotein (P-gp, also known as ABCB1 or MDR1), an ATP-binding cassette (ABC) transporter, is highly expressed at the luminal membrane of brain endothelial cells and serves as a key efflux pump at the blood brain barrier (BBB) [9, 10]. P-gp actively transports a broad range of chemotherapeutic agents and small molecules back into the bloodstream, thereby contributing to CNS drug resistance and poor therapeutic response in brain tumors, including GBM [11, 12]. Anthracyclines (E.g. doxorubicin, doxil), vinca alkaloids (E.g. vincristine, vinblastine) and taxanes (E.g. paclitaxel) are all common chemotherapy substrates of P-gp that are actively pumped out of tumor cells [13]. Increased activity of P-gp in both tumor-associated endothelium and residual BBB-protected regions limits drug accumulation in tumor parenchyma which contributes to the failure of many CNS-directed therapies [14, 15]. Previous studies have investigated the inhibition of P-gp as a strategy to overcome chemoresistance in glioma and GBM. For instance, agents such as Ibrutinib and Loperamide have been shown to enhance the intracellular accumulation of chemotherapeutic drugs in glioma models by inhibiting P-gp-mediated efflux [16, 17].

Angulin-1, also referred to as the lipolysis-stimulated lipoprotein receptor (LSR), is highly expressed in both endothelial and epithelial tissues, where it contributes to barrier function maintenance [7, 8]. Genetic deletion of LSR in mice results in embryonic lethality, although previous studies show CNS-specific loss does not affect bicellular TJ protein localization or permit leakage of large endogenous molecules into the brain parenchyma [18]. Interestingly, in disease models characterized by BBB disruption, such as multiple sclerosis and ischemic stroke, decreased LSR expression correlates with regions of heightened inflammation and vascular permeability [8]. Beyond the endothelium, LSR is also enriched in neural subpopulations, including glial cells, neurons, and myelin membranes, suggesting a role in cholesterol trafficking and neural homeostasis [19].

Pharmacologic targeting of Angulin-1/LSR has demonstrated therapeutic potential.

Specifically in metastatic lung cancer mouse models, Angubindin-1, a *Clostridium perfringens* iota toxin-derived peptide that binds tightly to Angulin-1/LSR, facilitated rapid CNS delivery of a 5.3 kDa antisense oligonucleotide (ASO), targeting Metastasis-Associated Lung Adenocarcinoma Transcript 1 (MALAT1). A significant increase in ASO concentration within the brain and spine of these animals was observed within 2 hours of administration at low doses (10–30 mg/kg); indicating effective modulation of endothelial permeability via Angulin-1/LSR inhibition [20].

Beyond the CNS, elevated Angulin-1/LSR expression has been observed in several non-CNS solid tumors, including ovarian and endometrial cancers, where it correlates with advanced disease stage and poorer prognosis [8, 21-23]. In rodent models of gynecologic cancers, therapeutic targeting of angulin-1 with monoclonal antibodies resulted in reduced tumor growth and extended survival [21]. This upregulation is hypothesized to support cholesterol uptake and nutrient trafficking in cancer cells, contributing to enhanced proliferation and invasiveness [24, 25]. No previous research on the role of angulin-1 inhibition on glioma nor correlative tumor endothelium has been performed to date.

Based on these previous findings, we hypothesize that high angulin-1/LSR expression contributes to both BBB integrity and glioma aggressiveness through its role in maintaining tricellular TJ architecture and modulating tumor-endothelial interactions. Here, we investigate the effects of angulin-1/LSR inhibition using angubindin-1, first on brain endothelial tight junctions, then on glioma cell progression, and finally in combination with the cytotoxic, P-gp substrate, Doxil in rodent glioma models. These studies aim to elucidate the dual role of angulin-1/LSR in regulating blood-tumor barrier permeability and promoting glioma progression, with implications for overcoming P-gp-mediated drug resistance at the BBB.

## Materials and Methods

### Chemicals and Reagents

Dulbecco’s Modified Eagle Medium (DMEM) high glucose and Eagle’s Minimum Essential Medium (EMEM) were obtained from Invitrogen (Carlsbad, CA, USA). Penicillin streptomycin, Hanks’ Balanced Salt Solution (HBSS), phosphate-buffered saline (PBS), fetal bovine serum (FBS), and 10X trypsin-EDTA were purchased from Invitrogen (Carlsbad, CA, USA). The Pierce BCA Protein Assay Kit, B27 supplement minus vitamin A, N2 supplement, GlutaMAX supplement, and all other reagents were acquired from ThermoFisher Scientific (Wilmington, DE, USA).

### Cell Culture

Rat brain microvascular endothelial cells (RBMVEC) were obtained from Cell Applications (Cell Applications, San Diego, CA, USA) and cultured following the manufacturer’s protocol. S635 cells were gifted from the D. Bigner Lab (Tisch Brain Tumor Center, Duke University, Durham, NC, USA) and were cultured in DMEM high glucose media supplemented with 10% FBS and 1% penicillin streptomycin (10,000 U/mL). LN-229 cells were purchased from ATCC (Manassas, VA, USA) and grown in DMEM media supplemented with 5% FBS and 1% penicillin streptomycin and cultured following the manufacturer’s protocol. *ABCB1*-transfected MDR-19 cells were maintained in EMEM supplemented with 10% FBS, 1% penicillin– streptomycin, and 2 mg/mL G418. All cells were maintained at 37°C in a humidified 5% CO_2_ incubator and cultured following the manufacturer’s protocol.

### Assessment of P-glycoprotein Function and Expression via Rhodamine 123 Efflux and Flow Cytometry

To evaluate ABCB1 (P-glycoprotein, P-gp) transporter function, rhodamine 123 efflux assays were performed in MDR-19 cells, which overexpress human P-gp, and in S635 rat glioma cells. The assay was conducted with or without the P-gp inhibitor valspodar, following established protocols [12]. Briefly, cells were trypsinized, washed, and incubated in phenol red–free DMEM containing 0.5 µg/mL rhodamine 123 at 37 °C for 30 minutes. Treatments included vehicle control, valspodar (3 µg/mL), or Angubindin-1 (600 µg/mL). After incubation, the rhodamine-containing medium was replaced with fresh, rhodamine-free medium containing the corresponding treatments, and cells were incubated for an additional hour to allow efflux. Cells were washed with PBS and analyzed by flow cytometry to assess intracellular rhodamine fluorescence, with higher fluorescence indicating reduced P-gp activity.

To assess P-gp expression, flow cytometry was performed on MDR-19 and LN-229 human glioma cells using a PE-conjugated anti-human CD243/MDR-1 antibody (UIC2 clone, #348608, BioLegend, San Diego, CA, USA) or a PE mouse IgG2a κ isotype control (#400214, BioLegend, San Diego, CA, USA). Cells were plated in triplicate in 96-well plates, centrifuged at 750 rpm for 5 minutes, and incubated with 200 µL of 2% BSA blocking buffer containing 5 µL of antibody, isotype control, or blocking buffer alone. Plates were wrapped in foil and incubated for 20 minutes at room temperature, followed by two PBS washes. Samples were transferred to flow cytometry tubes for analysis. Data acquisition and analysis were performed using FlowJo software (version 10.6.1; Tree Star, Inc., Ashland, OR, USA).

### Protein Synthesis against Angulin-1/LSR, Negative Control, Mutation of Angulin-1/LSR Binding Site

Angubindin-1, a clostridium iota toxin previously shown to bind to Angulin-1/LSR, was created by the method of expression construct generation: 27282-X01-566 His6-MBP-tev-angubindin-1 optEc.12. The template for 27282-T01 tev-angubindin-1 optEc was ordered from ATUM and reverse BP’d via Gateway Recombinational Cloning to make Entry clone 27282-E01. Entry clone was then cloned via Gateway Recombinational Cloning into bacterial vector pDest-566 to make Expression clone 27282-X01-566 His6-MBP-tev-angubindin-1 optEc. Expression clone DNA was transformed into DH10B e. coli cells and grown in small culture and miniprepped (QIAGEN, Germantown, MD, USA) before being verified on an agarose gel. Small aliquots of confirmed clone DNA were transferred to the Core Expression Group. For bacterial expression, 50 mL seed culture (in MDAG medium) was grown overnight in 250 mL baffled flask at 37 °C and agitated at 250 rpm. Then 40 mL of the seed culture was added to 2 L of dynamite broth in New Brunswick BioFlow 110 fermenter (3 L tank capacity). The tank was aerated with pure oxygen at an impeller speed of 481 rpm to maintain dissolved oxygen at >50% and incubated at 37 °C. When A600 reached ∼7, IPTG was added at 0.5 mM, the tank was cooled to 16°C, and growth continued overnight (19 hour after temperature change). Following growth, E. coli cells were harvested by centrifuging at 6328 x g at 4 °C for 20 min and cell pellets frozen at -80 °C before further processing. Pellet lysis was performed from cell pellet two-liter fermentation (76.8 gram pellet) and was resuspended in 396 mL of 20 mM HEPES, pH 7.4, 300 mM NaCl, 1 mM TCEP 1:100 (v:v) Protease Inhibitor Cocktail (Sigma-Aldrich, St. Louis, MO, USA) and lysed by passing through a microfluidizer twice at 10,000 psi. The lysate was clarified by ultracentrifugation at 8,000 x g at 4°C for 90 minutes. The cleared lysate was filtered through a 0.45 μm filter, and imidazole added to a final concentration of 35 mM. All purification steps were performed on a Bio-Rad NGC chromatography system at room temperature. Special steps were included to reduce endotoxin levels. All buffers were 0.2 mm filtered, stored in refrigerator, and used on ice to keep as cold as possible. A 50-ml HisTrap HP column was equilibrated in 10 column volumes (CV) 93% Buffer A (20 mM HEPES pH 7.4, 300 mM NaCl, 1 mM TCEP, 50 mM imidazole) and 7% Buffer B (Buffer A with 500 mM imidazole). Cleared lysate sample was loaded onto the column followed by 30 CV of Buffer C (Buffer A with 0.1% Trition X-114) using ice cold buffer. Column was further washed with 30 CV of ice-cold Buffer A. Protein was eluted from the column with 10 CV gradient from 0-100% Buffer B (0-500 mM imidazole). Fractions were collected across the gradient and analyzed by SDS-PAGE/Coomassie Blue staining. Elution fractions containing angubindin-1 were pooled. Pooled sample was TEV digested with 2.5% v/v addition of TEV protease and dialyzed into Buffer A using 10 kDa molecular weight cut off SnakeSkin dialysis tubing. TEV protease produced in house using similar endotoxin lowering protocol to maintain low endotoxin level of sample. A 50 mL HisTrap HP column was equilibrated in 10 CV Buffer A. TEV digested and dialyzed sample was loaded onto the column and washed with 10 CV of Buffer A. The column was eluted in a bump to 100% B. Fractions were collected throughout the load and wash and analyzed by SDS-PAGE/Coomassie Blue staining. Elution fractions containing Angubindin-1 were pooled. Pooled protein was concentrated using an Amicon stirred-cell concentrator. Final analysis was completed included endotoxin analysis, SDS-PAGE/Coomassie stained gel, and protein concentration.

### Mass Spectrometry (LC/MS), Circular Dichroism, Dynamic Lighting Scattering, Differential Scanning and Fluorimetry, and Mass Photometry

Found in “Supplementary Materials and Methods” section.

### Cellular Fractionation and Localization of Angulin-1/LSR in RBMVEC

To determine the subcellular localization of Angulin-1/LSR in RBMVEC, the Cell Signaling Cellular Fractionation Kit (Cell Signaling Technology, Danvers, MA, USA) was used to separate cells into membrane/organelle, cytoplasmic, and nuclear/cytoskeletal fractions. Approximately 5 × 10^6^ RBMVEC were seeded in each T-75 flask and cultured for 1, 2, 4, 8, 24 hours. Cells were then washed with cold PBS, trypsinized, and neutralized using growth medium to inactivate trypsin. The cell suspension was centrifuged at 220 × g for 5 minutes. After aspiration of the supernatant, the cell pellet was resuspended in 500 µL of cold PBS and processed according to the manufacturer’s protocol for fractionation. The resulting fractions were subsequently analyzed by SDS-PAGE followed by immunoblotting. Membranous Apoptosis-Inducing Factor (AIF) (Cell Signaling Technology, Danvers, MA, USA) was used as a loading control for membrane fractions, mitogen-activated protein kinase (MEK1/2) (Cell Signaling Technology, Danvers, MA, USA) for cytoplasmic fractions, and Histone H3 (Cell Signaling Technology, Danvers, MA, USA), or nuclear fractions enabling normalization of protein expression levels.

### Co-immunoprecipitation of Angulin-1/LSR and Angubindin-1 in RBMVEC and LN-229 Cells

Co-immunoprecipitation (Co-IP) was performed to investigate the interaction between Angulin-1/LSR and Angubindin-1. RBMVEC and LN-229 cells were seeded at a density of 5 × 10^6^ cells per T-175 flask and treated for 24 hours with either vehicle control, negative control of 10 µg/mL, or Angubindin-1 at concentrations of 75, 150, 300, and 600 µg/mL. Following treatment, cells were washed with PBS, trypsinized, and centrifuged at 300 × g to collect the cell pellet. Cell lysis was carried out using a buffer containing 20 mM Tris-HCl, 2 mM EDTA, 150 mM NaCl, and 0.2% Triton X-100, supplemented with a 1× Protease/PhosphataseArrest inhibitor cocktail (G-Biosciences, St. Louis, MO, USA). Protein concentration was quantified using the Pierce Protein Assay Kit (Thermo Fisher Scientific, Wilmington, DE, USA). Lysates were processed using the Pierce Co-Immunoprecipitation Kit (Thermo Fisher Scientific, Wilmington, DE, USA) according to the manufacturer’s instructions.

### Immunoblotting

Approximately, 2 × 10^6^ cells were plated per well in a T-75 flask and grown until confluent then treated with 10 µg/mL vehicle control, negative control, and 600 µg/mL of Angubindin-1 for 0, 1, 2, 4, 8, 24 hour. Cells were lysed with lysis buffer containing 20 mM Tris HCL, 2 mM EDTA, 150 mM NaCl, and 0.2% Triton-X100 with Protease-PhosphataseArrest [100X] cocktail (G-Biosciences, St. Louis, MO, USA). Protein concentration was determined using Pierce Protein Assay kit. Each sample contained 25µg of denatured proteins resolved on 4-20% Mini-Protean TGX precast gels (Bio-Rad, Hercules, CA, USA). Dry transfer method on PVDF membrane was used for protein transfer. Nonspecific proteins were blocked by incubating the membrane in 3% BSA blocking buffer for 1 hour at room temperature. 1:1000 dilution of primary antibodies Occludin (#ABIN500409), Claudin-5 (#ABIN6260869), ZO-1 (#ABIN675024), Tricellulin/MARVELD2 (#ABIN7264716), and Angulin-1/LSR (# ABIN6737870) were used and purchased from Antibodies Online (Limerick, PA, USA). Vinculin mAb (#4650), GAPDH mAb (#2118), HA-tag rabbit mAb (#3724), or DYKDDDDK Tag (D6W5B) Rabbit mAb (#14693) from Cell Signaling (Danvers, MA, USA) were diluted to 1:1000 concentration. Membranes were incubated with primary antibodies overnight at 4°C. The following day, the membranes were washed 3 times in 1X PBST for 5 mins. Secondary anti-rabbit IgG, HRP-linked antibody (#7074) were purchased from Cell Signaling (Danvers, MA, USA), and used for respective secondary antibody at 1:1000 concentration for 1 hour at 4°C. Membranes were washed again 3 times in 1X PBST for 5 min. Immunoblot images were detected using Bio-rad ChemiDoc Imaging System (Bio-rad, Hercules, CA, USA).

### Quantitative Reverse Transcription–Polymerase Chain Reaction

Total RNA was isolated from RBMVEC using the RNeasy Mini Kit (Qiagen, Germantown, MD, USA) according to the manufacturer’s protocol. For gene expression analysis, 0.5 μg of total RNA was reverse transcribed into cDNA using the Transcriptor Universal cDNA Master Kit (Sigma-Aldrich, St. Louis, MO, USA). For detection of oligonucleotides and quantification of gene expression, reverse transcription quantitative PCR (qRT-PCR) was performed using the High-Capacity cDNA Reverse Transcription Kit (ThermoFisher Scientific, Waltham, MA, USA) and run on a ViiA 7 Real-Time PCR System (Applied Biosystems). Gene expression levels were calculated using the 2^−ΔΔCt^ method. All qRT-PCR experiments were conducted in triplicate, with expression levels normalized to 18S ribosomal RNA (rRNA).

The primer and probe sets used for rat gene targets were as follows:

| Gene | Company | Assay ID |
| --- | --- | --- |
| LSR | Bio-Rad | qRnoCED0020018 |
| **Claudin-5** | Bio-Rad | qRnoCED000686 |
| **ZO-1** | Bio-Rad | qRnoCID0001801 |
| **MARVELD2 (Tricellulin)** | Bio-Rad | qRnoCED0018999 |
| **Occludin** | Bio-Rad | qRnoCED0011780 |

### Small Interfering RNA (siRNA) Knockdown of Angulin-1/LSR

RBMVEC were transfected with small interfering RNAs (siRNAs) targeting Angulin-1/LSR (50 nmol/L) or non-targeting control siRNAs (ThermoFisher Scientific; Wilmington, DE, USA) using Lipofectamine 2000 (Invitrogen; Carlsbad, CA, USA) following the manufacturer’s protocol. Briefly, 0.5 × 10^6^ cells were seeded and transfected in complete growth medium for 24 hours. Following transfection, knockdown efficiency was assessed via quantitative PCR; comparing siLSR to siControl cells. Cells demonstrating successful knockdown were used in subsequent experimental assays as described in the following sections.

### Orthotopic Glioma Implantation and Drug Treatment

All animal procedures were conducted in compliance with the National Institutes of Health Animal Use and Care (ACUC) Guidelines. Female Fischer 344 rats (5 weeks old) and immunodeficient Sprague Dawley Rag2/Il2rg (SRG) rats (5 weeks old) were purchased from Charles River Laboratories (Wilmington, MA, USA). Animals were anesthetized via inhalation of isoflurane (5% for induction and 2% for maintenance in oxygen) and positioned in a stereotactic head frame. A total of 1 × 10^4^ S635 or 1 × 10^6^ LN-229 GBM cells suspended in 3 or 10 μL of HBSS were stereotactically injected into the right hemisphere of the brain (coordinates: 2 mm anterior and 2 mm lateral to bregma; 4 mm depth) using a 28 or 33 gauge Hamilton syringe attached to a UMP3T microinjection pump (World Precision Instruments, Sarasota, FL, USA) at a controlled rate of 0.5-2 μL/min. Seven days post-implantation, rats were randomized into four treatment groups (*n* = 6-10 per group). Animals received weekly tail vein injections of angubindin-1 (10 mg/kg), followed by Doxil (liposomal doxorubicin, 3 mg/kg) administration after 1 hour of angubindin-1 treatment starting from 1 or 3 week of post-implantation (S635/Fisher and LN-229/SRG, respectively).

### Magnetic Resonance Imaging (MRI) and Analysis

MRI was conducted using a 4.7-Tesla Bruker Avance III MRI system equipped with ParaVision 6 software (Bruker, Billerica, MA, USA). Rats were anesthetized and secured on a custom-built cradle featuring a nose cone and bite bar to minimize head movement during imaging. Prior to scanning, a tail vein catheter was inserted and maintained with heparinized saline to allow intravenous contrast administration. T1-weighted images were acquired using Gadovist™ (0.1 mmol/kg) as the contrast agent. After acquiring localizer scans, axial slices of 1 mm thickness were obtained to cover the entire brain. MRI data were analyzed using ImageJ software (version 1.53; National Institutes of Health, Bethesda, MD, USA).

### Measurement of Doxil Biodistribution

Four hours following the final Doxil administration on day 14, animals were deeply anesthetized, and intracardiac blood was collected. To eliminate residual intravascular Doxil, transcardial perfusion was performed using saline. The tumor-bearing hemisphere and corresponding contralateral brain regions were then harvested, homogenized, and incubated in acidified ethanol at 4 °C for 24 hours to extract Doxil. Following extraction, samples were centrifuged at 16,000 × g for 25 minutes at 4 °C, and the resulting supernatant was collected for analysis. Doxil concentrations in brain tissue and plasma were quantified using a fluorometric assay on a Cytation 5 microplate reader (BioTek, VT, USA), as previously described [26].

### Histology

To assess the histological effects of each treatment, four animals from each group were sacrificed on day 14 post-tumor implantation. Animals were deeply anesthetized, and transcardial perfusion was performed using 4% paraformaldehyde (PFA) in PBS for fixation. Following perfusion, brains were extracted, immersed in 30% sucrose solution for cryoprotection, and subsequently embedded in OCT compound. Coronal brain sections were prepared at a thickness of 10 μm using a cryostat (Leica Biosystems, Buffalo Grove, IL, USA). Tissue sections were mounted on glass slides and stained with hematoxylin and eosin (H&E) using a commercial staining kit (Abcam, Cambridge, MA, USA) according to the manufacturer’s protocol. Whole-slide imaging was performed using a fluorescent stereo microscope (M165 FC, Leica Biosystems

### Immunohistochemistry

Rat brains were 4% paraformaldehyde-fixed, embedded in paraffin and sectioned at 5 μm thickness. Routine hematoxylin and eosin (H&E) staining were performed using the Sakura® Tissue-Tek® Prisma™ automated stainer (Sakura Finetek USA, Inc., Torrance, CA, USA). The slides were cover slipped using the Sakura® Tissue-Tek™Glass® automatic cover slipper (Sakura Finetek USA, Inc., Torrance, CA, USA). Whole slide images were obtained at high resolution, 20X magnification, using an AT2 scanner (Aperio, Leica Biosystems, Buffalo Grove, IL, USA). Single staining for Angulin-1/LSR, Claudin-5 and ZO-1 and double staining for CD31/Angulin, and were performed on the Leica Biosystems Bond RX autostainer using the Bond Polymer Refine Kit (Leica Biosystems, Buffalo Grove, IL, USA, #DS9800), with omission of the PostPrimary reagent, DAB and Hematoxylin. After antigen retrieval with EDTA (Bond Epitope Retrieval 2), sections for single staining were incubated for 30’ followed by Leica polymer reagent kit, secondary antibody Biotinylated Horse anti-mouse, streptavidin HRP, OPAL 690 Fluorophore, and OPAL Fluorophore 520 (AKOYA Biosciences, Marlborough, MA, USA). CD31 antibody complex was heat stripped with EDTA, sections were then incubated 60’ with Angulin-1, followed by OPAL fluor 690, and slides. Sections were removed from the bond, DAPI stained, and coverslipped with Prolong Gold AntiFade Reagent (Thermo Fisher Scientific, Waltham, MA, USA). For the double stain, after antigen retrieval with EDTA (Bond Epitope Retrieval 2) sections were incubated for 60 minutes with CD31 (Abcam, Waltham, MA, USA) followed by Leica polymer reagent kit and OPAL Fluorophore 520 (AKOYA Biosciences, Marlborough, MA, USA). Sections were then blocked with horse serum and incubated for 30 minutes. Sections were removed from the Bond, DAPI stained, and coverslipped with Prolong Gold AntiFade Reagent. Images were captured using the Aperio Scanscope FL whole slide scanner (Leica Biosystems, Buffalo Grove, IL, USA) into digital images. All image analysis was performed using Halo imaging analysis software (v3.3.2541.423, Indica Labs, Corrales, NM, USA) and image annotations were performed. Small and large vessels within 5-14 randomly selected areas (0.568 mm^2^) were evaluated for the presence and absence of CD31 and Claudin-5. IMARIS Surface analysis was generated for each image in which Machine Learning Segmentation was trained and applied for each individual junctional marker followed by the intensity sum was collected, based on each marker’s fluorescent expression.

### Transcriptomic Data Acquisition and Processing/Bulk RNA-sequencing analysis

Transcriptomic and clinical data were obtained from two publicly available sources: the Pediatric Brain Tumor Atlas (PBTA) and The Cancer Genome Atlas (TCGA). For TCGA, the GBM multiforme (TCGA-GBM) cohort (n = 157) was accessed, and both clinical metadata and RNA-sequencing (RNA-seq) data were retrieved using the TCGA biolinks R package [27]. Only high-grade glioma (HGG) samples were retained for analysis. For PBTA, bulk RNA-seq data were downloaded from the PedcBioPortal, with genomic annotations provided by the Children’s Brain Tumor Network (CBTN), the Pacific Neuro-Oncology Consortium (PNOC), and partners via the Gabriella Miller Kids First Data Resource Center. To focus on HGG, samples categorized under the histological group “Diffuse astrocytic and oligodendroglial tumor” (*n* = 676) and with available RNA-seq data were initially selected. Samples labeled as “WXS” or “Serum” were excluded to ensure that LSR expression measurements originated from brain tumor tissues. Additionally, samples from recurrent tumors or deceased patients were removed. This filtering yielded a final cohort of 259 PBTA samples. To integrate data across the PBTA and TCGA cohorts and correct for potential batch effects, log-transformed expression data were adjusted using the ComBat function from the sva R package, with TCGA used as the reference distribution. Principal component analysis (PCA) was conducted to assess data structure before and after batch correction, and one outlier was removed from the PBTA cohort for downstream analysis (Supplementary Figure 2). Angulin-1/LSR expression was compared across subgroups defined by age group using boxplots and statistical testing via the Wilcoxon rank-sum test. Associations between continuous variables were evaluated using Pearson correlation, and results were visualized with scatter plots.

### Single-Cell RNA-Sequencing Analysis

Previously published single-nucleus RNA-sequencing (snRNA-seq) data from a cohort of 24 patients with histone H3 lysine 27-to-methionine (H3K27M) mutant diffuse midline glioma (DMG) was accessed [28]. One sample with a spinal cord primary tumor was excluded, and the analysis was restricted to frozen tissue samples processed using SMART-seq2 single-nucleus RNA-seq (snRNA-seq), yielding a final cohort of n=21 samples. Data analysis was performed using the Seurat R package (v5.0). Raw count matrices were log-normalized using the log1p function, and a total of 3,293 cells were retained for downstream clustering. Batch correction across samples was implemented using Harmony (v1.2.0), and Harmony-corrected embeddings were subsequently used to generate uniform manifold approximation and projection (UMAP) plots. Cell clusters were identified using the FindClusters function in Seurat and annotated based on prior classification in the original publication of the snRNA-seq data [28]. LSR gene expression data were extracted using Seurat’s GetAssayData function. The processed data supporting this analysis are provided in Supplemental data. For access to the raw snRNA-seq files, refer to the original publication [28].

### Statistical Analysis

All results are presented as mean ± standard error of the mean (SEM) from experiments performed in triplicate (n = 3). Statistical significance between groups was assessed using unpaired t-tests or one-way analysis of variance (ANOVA) followed by Dunnett’s multiple comparison test for post hoc analysis. A p-value of less than 0.05 was considered statistically significant. All analyses were conducted using GraphPad Prism software (version 9.3; GraphPad Software, San Diego, CA, USA).

## Results

### Angulin-1/LSR is highly expressed on brain endothelium and localized to the cellular membrane

To assess the cellular distribution of Angulin-1/LSR expression, we examined single-nucleus RNA sequencing (snRNA-seq) data from a previously published dataset comprising 21 patients with DMG (3,293 nuclei and 13 distinct clusters) [28] (Figure 1A). Angulin-1/LSR expression was detected variably across identified cell populations (Figures 1A). Among non-tumor clusters, endothelial and myeloid cells exhibited the highest Angulin-1/LSR expression, while T-cells displayed low expression levels (Figure 1A). Within tumor cell clusters, Angulin-1/LSR expression was primarily localized to AC-like (astrocyte like) and S-like (S cell cycle phase) subtypes, with minimal or absent expression observed in other malignant cell populations (Supplementary Figure 1). To further explore the relationship between Angulin-1/LSR expression and patient age, we analyzed publicly available bulk RNA-sequencing datasets from 413 patients with high-grade glioma (HGG) using harmonized data from the TCGA and PBTA datasets, applying batch correction (Supplementary Figures 2-4). Angulin-1/LSR was expressed across different age groups without significant differences among patients under 18 years, 19–39 years, and 40 years and older (Supplementary Figure 5). Additionally, no significant correlation between Angulin-1/LSR expression and age was observed, as shown by scatterplot analysis (Supplementary Figure 6-8).

**Figure 1.**
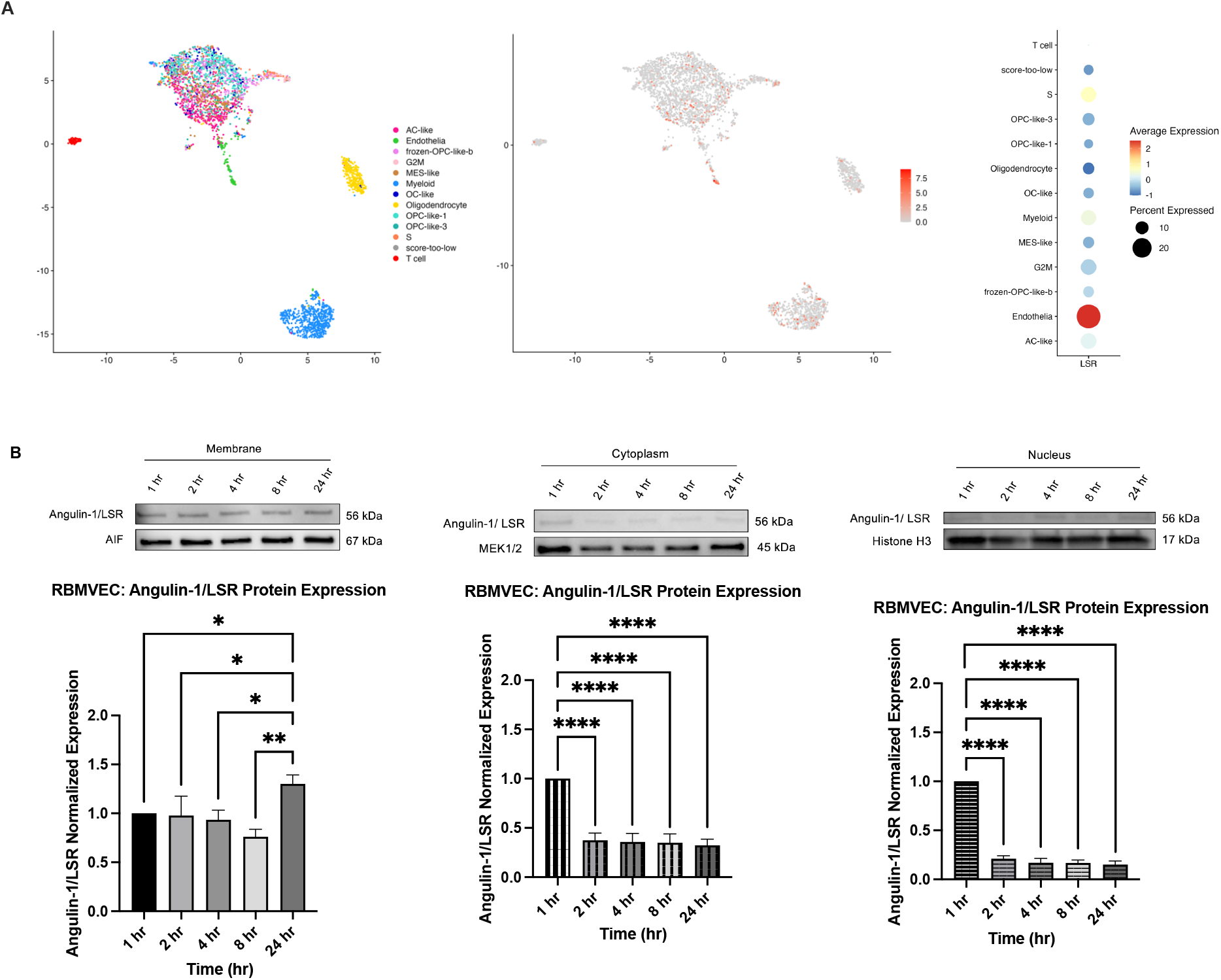
A-B. Angulin-1/LSR expression in high-grade glioma cell types and minimal change over time non-treated RBMVEC. (A) UMAP plot showing the cellular distribution of Angulin-1/LSR expression across identified clusters. Expression values were log-transformed using the *log1p()* function to normalize data and account for zero values. Points are jittered and colored according to cell type annotation. Dot plot summarizing the proportion and average expression of Angulin-1/LSR across cell clusters. Dot size represents the percentage of cells expressing the gene, while color intensity reflects the mean expression level per cluster. (B) Representative immunoblots showing Angulin-1/LSR protein levels in RBMVECs over a 24-hour period under untreated conditions. Quantification of Angulin-1/LSR expression by densitometry, normalized to the AIF, MEK1/2, or Histone H3 membrane loading control. Quantification of Angulin-1/LSR expression by densitometry, normalized to the AIF, MEK1/2, or Histone H3 membrane loading control. Data represent the mean ± SEM from three independent biological replicates (*n* = 3). Statistical comparisons were made across all time points (1, 2, 4, 8, and 24 hours). \**p* < 0.05, \*\**p* < 0.001, \*\*\*\**p* < 0.00001 indicate significant differences relative to other time points.

To confirm previous studies showcasing membrane cellular expression of Angulin-1/LSR, we first evaluated Angulin-1/LSR localization on rat brain endothelium [29]. Our studies demonstrated Angulin-1/LSR is predominantly localized on the membrane of untreated RBMVEC (confirmed with loading control of mitochondrial membrane protein, AIF). We also noted minimal translocation to the cytoplasm or nucleus over 24 hours of growth after plating (Figure 1B). Quantitative analysis revealed no difference in Angulin-1/LSR expression from 1-8 hours after plating. Yet, expression levels significantly increased above baseline by 24 hours (*p* < 0.01).Cytoplasmic and nuclear fractions showed a significant decrease to more than half of baseline levels of Angulin-1/LSR expression from 1-24 hours (*p* < 0.00001), with confirmatory loading control proteins of MEK1/2 and histone H3, respectively. LSR/Angulin-1 exhibits dynamic regulation at the endothelial cell membrane under basal conditions without intracellular redistribution. Our findings show that Angulin-1/LSR exhibited dynamic regulation at the endothelial cell membrane under basal conditions without intracellular redistribution.

### Angubindin-1 Modulates Angulin-1/LSR Protein–Protein Interactions in a Cell Type Dependent Manner and dimerizes at high concentrations

Using a clostridium iota toxin previously shown to bind to Angulin-1/LSR, His6-MBP-tev-angubindin-1 (angubindin-1) 3X-FLAG tagged, we aimed to evaluate the effect of protein inhibition on cellular and subcellular activity. Structural prediction using AlphaFold identified potential binding interfaces between Angubindin-1 and Angulin-1/LSR, including the four residue pairings of Arg531–Glu151, Tyr535–Lys176, Thr178–Tyr42, and Lys127–Asp49, suggesting a specific and multivalent interaction profile. These findings allowed for purity and integrity of Angubindin-1 creation (Supplemental Figure 9). Specifically, protein interactions between Angubindin-1 (75–600 µg/mL) and Angulin-1/LSR were assessed using co-immunoprecipitation in both endothelial and human glioma cells (LN-229). In RBMVECs, Angubindin-1 induced a dose-dependent increase in Angulin-1/LSR interaction, with statistically significant elevations observed at 300 µg/mL (*p* <0.01) and 600 µg/mL (*p* <0.001) compared to control (Figure 2A). In contrast, in LN-229, Angubindin-1 treatment resulted in a statistically significant decrease in Angulin-1/LSR interaction across the entire concentration range compared to control (Figure 2B). These findings suggest that Angubindin-1 regulates Angulin-1/LSR in endothelium by dose-dependently increasing interaction of this tricellular protein, yet in glioma cells, it dose-independently decreases interaction. Based on this data, subsequent *in vitro* studies utilized the 600 µg/mL dosing to evaluate downstream effects on the role of Angubindin-1 to modulate endothelial membrane junctional expression versus the cell-to-cell interactions in aggressive/infiltrative glial cells (Supplementary Figures 9-14, Supplementary Table 1). aggressive/infiltrative glial cells.

**Figure 2.**
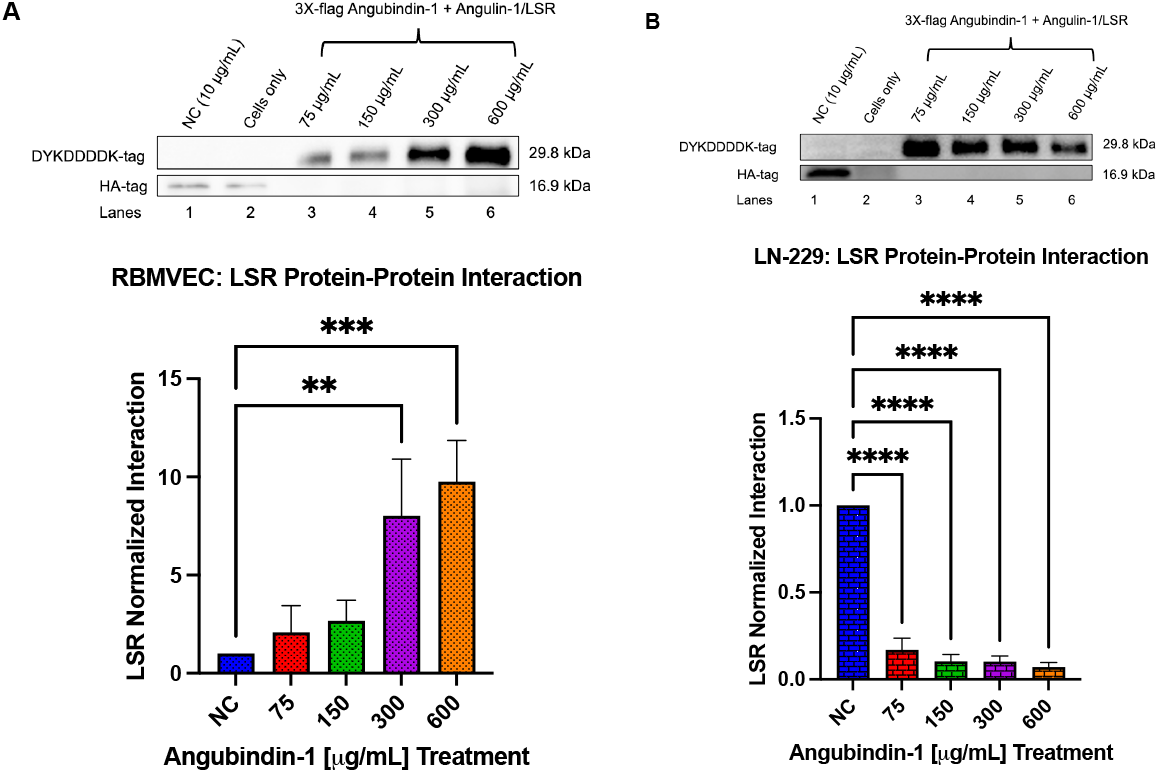
A–B. Angubindin-1 dose-dependently increased Angulin-1/LSR Protein–Protein Interactions in endothelium only. Cells were treated with Angubindin-1 at increasing concentrations (75–600 µg/mL) for 24 hours, followed by co-immunoprecipitation using 6 µg/mL anti-LSR antibody to evaluate Angulin-1/LSR interactions. (A, B) Representative immunoblots of 3X-FLAG-Angubindin-1 and Angulin-1/LSR in RBMVEC and LN-229 cells, respectively. Densitometric quantification of Angulin-1/LSR interactions, normalized to the HA-tagged negative control (NC) loading control. Angubindin-1 elicited a dose-dependent effect on Angulin-1/LSR expression: increased expression was observed in RBMVEC, while a dose-dependent decrease was seen in LN-229 glioma cells. All experiments were conducted in triplicate (*n* = 3) across three independent biological replicates. Data are presented as mean ± SEM. Statistical comparisons between negative control (vehicle control) and Angubindin-1-treated groups were analyzed using one-way ANOVA followed by Dunnett’s post hoc test. \*\**p* < 0.01, \*\*\**p* < 0.001, \*\*\*\**p* < 0.0001.

### Angubindin-1 Disrupts Bicellular and Tricellular Tight Junction (TJ) Integrity and LSR Inhibition Downregulates Junctional mRNA Expression

To evaluate the effect of Angubindin-1 on tight junction regulation, we assessed both protein and mRNA expression levels of bicellular (Occludin, Claudin-5, ZO-1) and tricellular (Tricellulin/MARVELD2, Angulin-1/LSR) tight junction components in RBMVECs, along with Angulin-1/LSR expression in LN-229 human glioma cells. Cells were treated with 600 µg/mL Angubindin-1 for up to 24 hours. We found that while occludin has a published half-life of approximately 6.2 hours in endothelial cells [30], angubindin-1 significantly reduced protein expression 2 and 8 hours post-treatment with a return to baseline by 24 hours (*p* < 0.05; Figure 3A). Claudin-5, which has a reported half-life ranging from 1.16 to 13.82 hours [30], exhibited a transient decrease at 2-24 hours after treatment (*p* < 0.05; Figure 3B). Similarly, ZO-1 protein expression, with a half-life of 5.1 to 7.7 hours [31], was significantly reduced between 1-24 hours (*p* < 0.05; Figure 3C). Evaluating tricellular junctions, angubindin-1 treatment demonstrated a significant decrease in Tricellulin/MARVELD2 expression, (un-published half-life) 1-8 hours later (*p* < 0.01), with a slight increase above baseline observed at 24 hours *p* < 0.05; Figure 3D). Angulin-1/LSR expression followed a similar biphasic pattern, showing significant decreases at 1, 2, and 4 hours (*p* < 0.01, *p* < 0.01, *p* < 0.001), a return to baseline at 8 hours, and a robust increase above baseline at 24 hours (*p* < 0.0001, Figure 3E). The half-life of Angulin-1/LSR has not been reported. LN-229 cells showed a consistent and significant reduction in Angulin-1/LSR expression after 1, 2, and 4 hours post Angubindin-1 (*p* < 0.001, *p* < 0.001, *p* < 0.01), with a subsequent increase above baseline at 24 hours (*p* < 0.0001, Figure 3F). Additionally, we assessed the impact of mutant-Angubindin-1 on the expression of tight junction proteins Claudin-5 and Angulin-1/LSR over a 24-hour time course. Treatment with 600 µg/mL mutant-Angubindin-1 resulted in a time-dependent reduction in Claudin-5 expression, with levels decreasing significantly at 2 hours (*p* < 0.0001) before gradually returning to near-baseline by 24 hours (Supplemental figure 15). In contrast, Angulin-1/LSR protein levels showed a sustained decrease over time, becoming significantly reduced by 24 hours (*p* < 0.00001). Analysis of mRNA expression in RBMVECs revealed downregulation of both bicellular and tricellular tight junction components post siLSR transfection of brain endothelium. Specifically, mRNA levels of Occludin, Claudin-5, and ZO-1 were significantly reduced compared to siControl transfected cells (*p* < 0.0001, *p* < 0.001, *p* < 0.01, respectively). Tricellular tight junction mRNAs, Tricellulin/MARVELD2 and Angulin-1/LSR also showed significant downregulation in siLSR endothelium (*p* < 0.0001, Figure 3G). Collectively, this mRNA expression data supported our decreased protein expression findings post Angulin-1/LSR inhibition from Angubindin-1.

**Figure 3.**
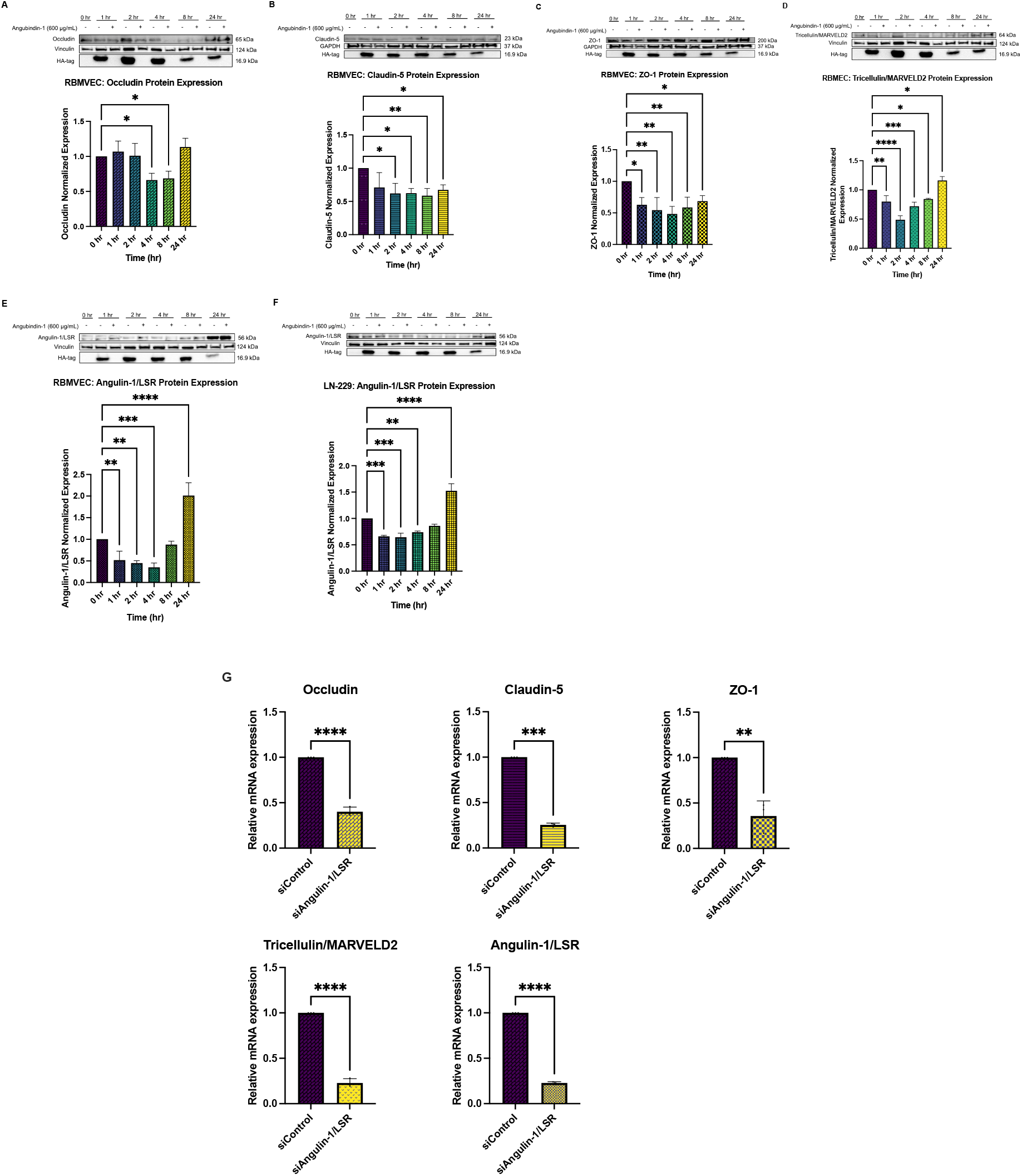
A–G. Angubindin-1 Decreases Bicellular and Tricellular Tight Junction Protein Component Expression in RBMVEC and Human Glioma Cells. (A-F) Protein expression levels of bicellular (Occludin, Claudin-5, ZO-1) and tricellular (Tricellulin/MARVELD2, Angulin-1/LSR) tight junctions were assessed by immunoblotting and densitometric analysis across 0, 1, 2, 4, 8 and 24 hours following treatment with 600 µg/mL Angubindin-1. Angubindin-1 induced time-dependent reductions in tight junction protein levels across the 24 hour time course. Only in tricellular junctions was an increase in expression above baseline seen at 24 hours. (G) The mRNA expression of both bicellular and tricellular junctions were evaluated after siLSR compared to siControl also demonstrated decreased levels of Occludin, Claudin-5, ZO-1, Tricellulin/MARVELD2, and Angulin-1/LSR. Graphs represent mean ± SEM from three independent biological replicates (*n* = 3). Statistical significance was determined using one-way ANOVA followed by Dunnett’s post hoc test. \**p* < 0.05, \*\**p* < 0.01, \*\*\**p* < 0.001, \*\*\**p < 0*.*0001*.

### Suppression of P-Glycoprotein (P-gp/ABCB1) Efflux Function by Angubindin-1 in Rodent Glioma Cells

P-glycoprotein (P-gp/ABCB1) is a key efflux transporter at the blood-brain barrier that restricts drug penetration into cells, thereby limiting therapeutic efficacy. Historically, P-gp has demonstrated close association and coordinated work with multiple TJs to limit CNS permeability[10]. To specifically assess Angubindin-1’s effect on P-gp function, we examined both S635 rat glioma cells as well as the human-derived glioma cells LN-229. Treatment with Angubindin-1 (600 µg/mL) significantly inhibited P-gp function in S635 rat glioma cells, as evidenced by increased intracellular accumulation of rhodamine 123 (a P-gp substrate) and compared to known P-gp inhibitor valspodar, rhodamine only and cells without fluorescent dye (Figure 4A). These results indicate that Angubindin-1 impairs P-gp–mediated efflux, thereby enhancing substrate retention in glioma cells. In contrast, the human glioma cell line LN-229 displayed low expression of P-gp as evidenced by UIC expression not being significantly different than isotype control and lower fluorescence expression compared with MDR-19 P-gp overexpressing cells (Figure 4B).

**Figure 4.**
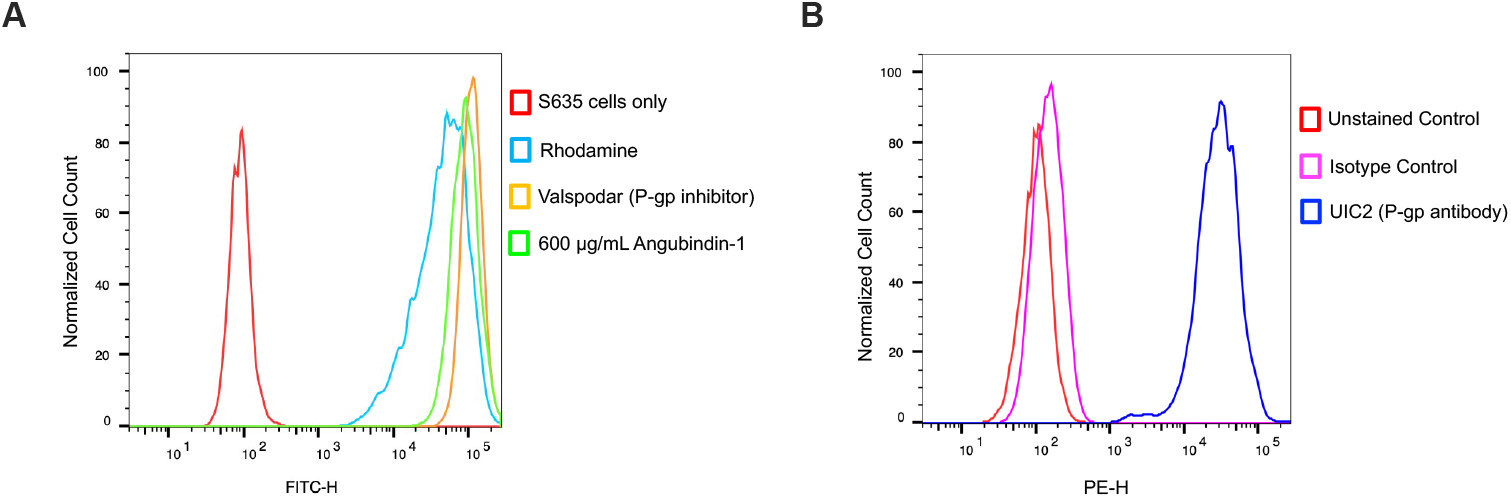
A-B. Angubindin-1 Inhibits P-glycoprotein (P-gp) Transporter Function and Confirms Low P-gp Expression in LN-229 Cells. (A) Treatment with Angubindin-1 increased intracellular accumulation of rhodamine 123, as indicated by a rightward shift in FITC-H fluorescence intensity, comparable to cells treated with the known P-gp inhibitor valspodar. This shift reflects inhibition of P-gp efflux activity. (B) P-gp expression was assessed using UIC2 antibody binding in MDR-19–transfected cells versus untreated LN-229 glioma cells. LN-229 cells showed no significant rightward shift in PE-H fluorescence, indicating low or absent P-gp expression, in contrast to P-gp–positive MDR-19 cells. Data was acquired and analyzed from three independent biological replicates (*n* = 3) using FlowJo software. Fluorescence intensity was used as an indicator of intracellular rhodamine retention and P-gp transporter function.

### Combination Angubindin-1 and Doxil Therapy Reduces Tumor Burden and Prolongs Survival in Immunocompetent Rats but Not in Human Glioma Xenografts

To assess the *in vivo* efficacy of combined Angubindin-1 and Doxil therapy, we first performed intracranial injections of S635 rat glioma cells in immunocompetent rats. Seven days post-implantation, animals received intravenous administrations of Doxil (3 mg/kg) and Angubindin-1 (10 mg/kg) until they met clinical endpoint criteria for Doxil cortical concentrations, MRI evaluation, and survival studies (Figure 5A). In the S635 rat glioma model, combination therapy significantly enhanced Doxil accumulation in the tumor-bearing hemisphere compared to the contralateral side (12.7 ng/g vs. 10.5 ng/g; *p* < 0.005) (Figure 5B). MRI analysis demonstrated that the greatest tumor volume reduction occurred with combination therapy (approximately 7 mm^3^) compared to Doxil alone (40 mm^3^) or Angubindin-1 alone (9 mm^3^), with statistical significance observed (*p* < 0.05) (Figure 5C). Histological analysis of H&E-stained brain sections corroborated the MRI findings, showing notably reduced tumor burden in Angubindin-1–treated animals (Figure 5D). Yet, assessments of junctional expression (Claudin-5, ZO-1, LSR) 14 days after Angubindin-1 first dose, displayed no differences compared to control; suggesting that inhibition of junctions occurs earlier, as evidenced by *in vitro* studies, to influence disease (Supplemental Figure 15). Survival analysis revealed that Angubindin-1 (10 mg/kg) alone modestly prolonged median survival (20 days), while combination treatment further extended survival to 24.7 days, compared to Doxil alone (19 days) and vehicle control (16 days; *p* < 0.05) (Figure 5E). Remarkably, Angubindin-1 therapy alone significantly improved survival over control (*p* < 0.0001), prompting further evaluation in a human-derived glioma model.

**Figure 5.**
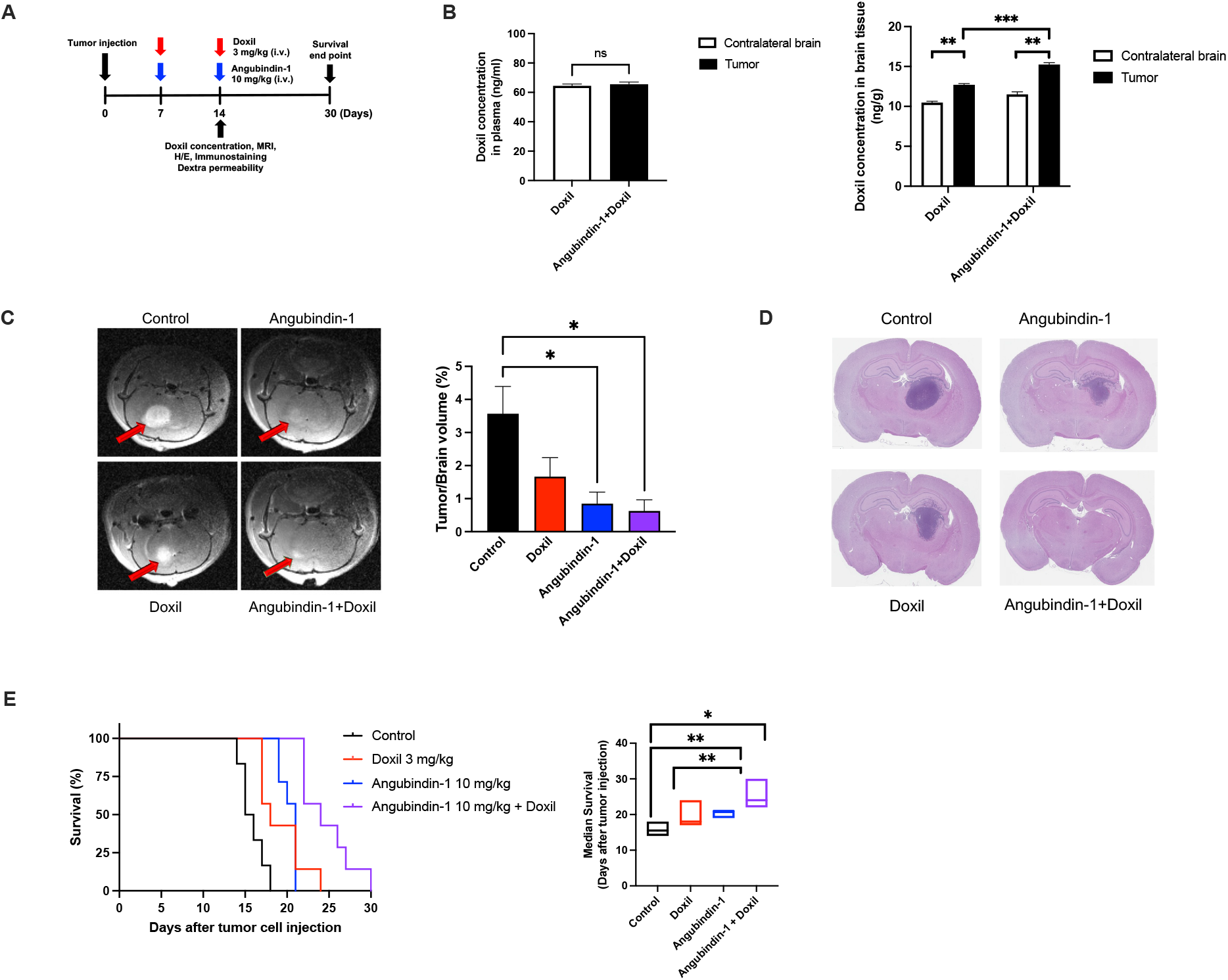
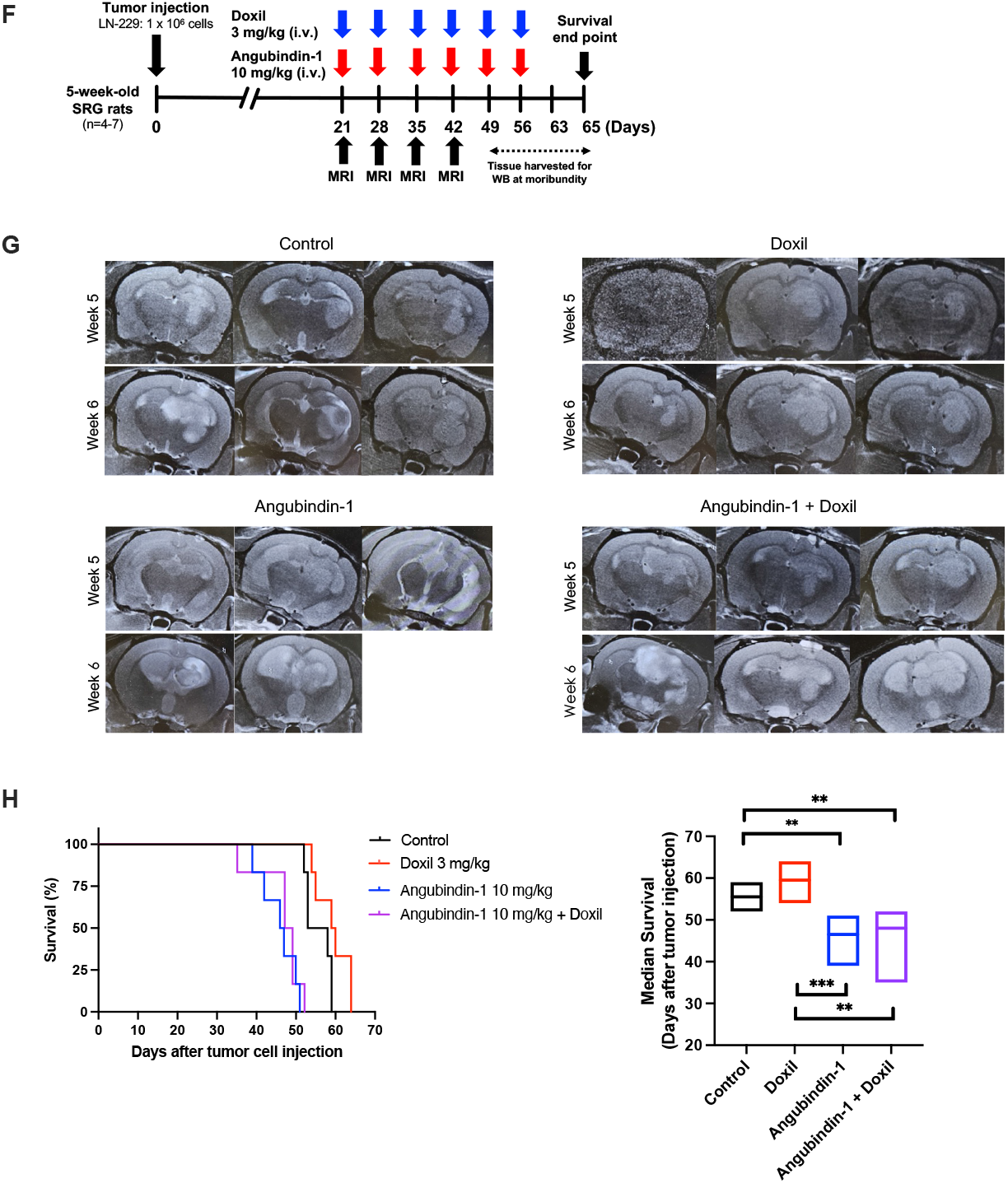
A–H. Combination Treatment with Angubindin-1 and Doxil Reduces Glioma Tumor Burden and Prolongs Survival in Preclinical Models. (A) Experimental timeline outlining treatment administration, MRI imaging, Doxil and Angubindin-1 blood/tissue concentrations, and survival monitoring in S635 rat glioma cells. (B) Co-administration of Angubindin-1 with Doxil did not alter plasma Doxil concentrations but significantly increased Doxil accumulation in the tumor-injected hemisphere (\**p* < 0.005). (C) Representative MRI images in the S635 rat glioma model show reduced tumor volume in the combination treatment group compared to vehicle, Angubindin-1 alone, or Doxil alone, with quantification confirming a significant reduction in tumor size (\**p* < 0.05). (D) H&E-stained brain sections reveal reduced tumor size in animals treated with Angubindin-1 alone or in combination with Doxil. (E) Kaplan–Meier survival curves demonstrate significantly prolonged survival in the combination therapy group (median survival: 24.7 days) versus control (16 days) (\**p* < 0.05, \*\**p* < 0.01). (F) Experimental timeline for LN-229 human glioma model. (G) MRI scans indicate that combination therapy is associated with additive tumor growth compared to vehicle control. (H) Kaplan–Meier analysis in the LN-229 model showed no significant survival advantage with either Angubindin-1 alone or in combination with Doxil.

To examine whether these effects translated to human-derived glioma cells, we utilized immunodeficient rodents (lacking B, T, and NK cells) intracranially injected with LN-229. Twenty one days post-implantation, animals received intravenous administrations of Doxil (3 mg/kg) and Angubindin-1 (10 mg/kg) until they met clinical endpoint criteria for Doxil cortical concentrations, MRI evaluation, and survival studies (Figure 5F)Under the same dosing regimen as S635 rat glioma cells, MRI revealed increased tumor growth in both Angubindin-1 alone and combination therapy groups relative to control (Figure 5G). In association, we observed a lower median survival in the animals treated with Angubindin-1 alone or in combination to Doxil. These findings suggested a potentially pro-proliferative effect of Angubindin-1effect in the immunocompromised setting in cells with low P-gp expression (Figure 5H).

## Discussion

GBM is the most aggressive and lethal primary brain tumor, characterized by rapid growth, diffuse infiltration, and resistance to therapy. The current standard of care includes maximal safe surgical resection followed by concurrent chemoradiation with temozolomide (TMZ), an FDA-approved alkylating agent [32]. Despite this multimodal approach, median survival remains approximately 18 months, with minimal improvement over the past two decades [33]. A major barrier to effective chemotherapeutic treatment is the blood–tumor barrier, which tightly regulates the endothelial interface and restricts drug penetration into the central nervous system. Although TMZ is BTB-permeable, its therapeutic efficacy remains modest, partly due to heterogeneous drug delivery and intrinsic tumor resistance mechanisms [34-36]. This underscores an urgent need for novel strategies that transiently and safely disrupt the BTB to enhance drug permeability without causing neurotoxicity.

In this study, we demonstrate the therapeutic potential of targeting brain endothelial tight junction proteins, such as Angulin-1/LSR, to transiently modulate BBB/BTB, inhibit P-glycoprotein (P-gp) efflux transporter activity, and enhance the delivery of the chemotherapeutic agent Doxil when combined with Angubindin-1 in rodent glioma models. These findings suggest a promising translational approach for addressing one of the key obstacles in GBM treatment: limited drug penetration across the BBB. RBMVEC widely used for modeling BBB function, provided a relevant *in vitro* system for studying endothelial barrier modulation [37]. The glioma models employed in this study include the S635 cell line, an IDH wild-type rat anaplastic astrocytoma derived from chemically induced tumors in Fischer 344 rats, and the LN-229 human glioma cell line, originally isolated from the right frontal parieto-occipital cortex [38], allowing us to explore both species, which can drive specific and translational discoveries towards BTB-targeted therapy.

Angubindin-1 has emerged as a promising modulator of the BBB/BTB, originally demonstrated to facilitate CNS delivery of antisense oligonucleotides by selectively targeting tricellular tight junctions [20, 39]. Building on these findings, our study aimed to evaluate Angubindin-1’s impact on Angulin-1/LSR expression and its capacity to enhance Doxil permeability across the BTB [20, 40, 41]. We first mined published single cell data exploring Angulin-1/LSR expression in well recognized pediatric glioma database (PBTA). Transcriptomic analyses revealed that Angulin-1/LSR expression is not age-dependent but is instead enriched in endothelial and myeloid cells, as well as in specific glioma cell subtypes (AC-like and S-like). These patterns support the idea that Angulin-1/LSR serves as a cell-type–specific therapeutic target, particularly in vascular-associated, treatment-resistant glioma populations, regardless of patient age [8, 29, 42]. We then confirmed that Angulin-1/LSR is dynamically regulated at the plasma membrane of brain endothelium, exhibiting a biphasic expression pattern with early downregulation and robust rebound at 24 hours; with minimal cytoplasmic or nuclear translocation. These minimal fluctuations likely represent intrinsic feedback mechanisms that preserve BBB homeostasis, as previously observed of TJ proteins responding to microenvironmental cues or cellular stress in other vascularized tumors or neurologic conditions [23, 43, 44]. Further, we demonstrated that Angubindin-1 is linked to Angulin-1/LSR interactions in a concentration, cell-type specific manner contrasting RBMVEC from human glioma cells. This dual behavior of Angulin-1/LSR inhibition and interaction suggests a context-specific effect on tricellular junction remodeling, possibly driven by differential expression of junctional scaffolding proteins or post-translational modifications [43-45]. This tunable activity offers a strategic opportunity to selectively disrupt the BTB in tumors while preserving normal endothelial barrier function.

At both the transcript and protein levels, Angubindin-1 induced transient downregulation of key bicellular (Occludin, Claudin-5, ZO-1) and tricellular (Tricellulin, Angulin-1/LSR) TJ components in RBMVECs, followed by delayed recovery, indicative of a reversible mechanism. This biphasic pattern mirrors the natural cycling of TJ assembly and disassembly during physiological regulation, yet with longer timing to allow for drug permeability [29, 45]. Additionally, we found that Angubindin-1 impaired P-gp function in rat glioma cells with high expression, increasing intracellular drug retention. The variable expression of P-gp across cell lines (higher in rat derived S635 than in human derived LN-229) aligns with known heterogeneity in glioma transporter profiles and explains differing treatment responses [46].

To enhance clinical relevance, we evaluated the therapeutic potential of Angubindin-1 alone and in combination with Doxil, the pegylated liposomal formulation of doxorubicin. Unlike free doxorubicin, which has limited ability to cross the BTB due to its high molecular weight and P-gp mediated efflux [47, 48], Doxil exhibited improved pharmacokinetics and prolonged circulation time, increasing the likelihood of passive accumulation in tumor tissue via the enhanced permeability and retention effect [49]. These properties make Doxil a more suitable candidate for BBB penetration in preclinical models and a rational choice for assessing the efficacy of barrier-modulating agents such as Angubindin-1, yet it still serves as a substrate for P-gp. *In vivo*, Angubindin-1 enhanced Doxil delivery and efficacy in immunocompetent rats bearing S635 glioma, significantly increasing intratumoral drug concentration, reducing tumor burden, and improving survival, alone and in combination with chemotherapy. These benefits are likely due to targeted disruption of Angulin-1/LSR-mediated bicellular and tricellular junctions and concurrent P-gp inhibition; consistent with earlier studies advocating transient TJ modulation to enhance chemotherapeutic access to brain tumors [16, 29, 46, 50, 51]. However, in immunodeficient rodents bearing LN-229 human glioma xenografts, Angubindin-1 failed to confer therapeutic benefit, likely due to low P-gp expression of these tumor cells, as well as the absence of an immune response at the site of tumor cell injection. These findings emphasize and support published work that the tumor microenvironment and BBB/BTB architecture, along with host immunity are critical determinants of therapeutic success [34, 52]. Altogether, our findings position Angubindin-1 as a potent, selective modulator of BBB and BTB integrity, capable of enhancing drug permeability through regulated disruption of tricellular junctions and efflux transporter inhibition; with a specific relationship to high P-gp expression tumor cells. Further work in patient-derived and immunocompetent glioma models will be essential to fully harness clinical potential of Angulin-1/LSR inhibition.

This study has several limitations that may impact interpretation and translational relevance. First, the therapeutic benefit of Angubindin-1 was observed only in the immunocompetent S635 rat glioma model, but not in the LN-229 human xenograft model, likely due to species-specific differences in BTB structure, drug transporter expression, and tumor immune microenvironment. The lack of an intact immune system in the xenograft model may have further limited our ability to assess immune–vascular interactions that influence barrier permeability [3, 53]. Second, while Angubindin-1 induced reversible disruption of tight junction proteins and enhanced drug delivery, we did not evaluate long-term safety or neurotoxicity associated with repeated barrier modulation. This is critical given prior reports of off-target effects and neuronal injury with barrier-disrupting agents [54]. Additionally, although we observed impaired P-glycoprotein (P-gp) function *in vitro*, and higher doxil brain concentrations with addition of angubindin-1 in S635 rodent models, the precise mechanism of transporter inhibition requires further validation with *in vivo* experiments and molecular characterization, which are quite difficult to assess in such dynamic brain tumor models. Finally, while Angulin-1/LSR expression was enriched in specific glioma subtypes and vascular-associated cells, we did not stratify treatment response based on expression levels. Given the known heterogeneity of GBM, future studies should assess whether Angulin-1/LSR can serve as a predictive biomarker for barrier-modulating therapies, potentially based on baseline expression of tumor cell P-gp and Angulin-1/LSR alike [42].

Our findings underscore the urgent need for adjunct therapies that can transiently and selectively modulate the BBB or BTB to improve drug delivery to malignant gliomas. Current chemotherapies often fail to achieve therapeutic concentrations within brain tumors due to the restrictive nature of the BBB/BTB, contributing to poor patient outcomes and treatment resistance [3, 35]. This study provides compelling evidence that Angubindin-1 modulates tight junction integrity, impairs efflux transporter function, and influences glioma progression by transiently increasing BBB/BTB permeability. Through glioma models, we demonstrated that Angubindin-1 facilitates enhanced chemotherapeutic delivery, specifically Doxil, into glioma tissue, supporting its role as a potential adjunctive agent in overcoming one of the most critical challenges in glioma treatment, limited drug penetration across the BTB. By targeting both bicellular and tricellular junctional components and/or by inhibiting P-gp, Angubindin-1 offers a novel and targeted approach to disrupt the BTB without causing permanent damage to normal vasculature; as evidenced by its transient effects. This represents an advancement in the development of strategies aimed at improving therapeutic access to the brain for aggressive diseases/conditions. Given the poor prognosis associated with high-grade gliomas and the limitations of current therapies, our findings motivate further research of BTB-modulating agents that can be safely integrated with standard or emerging treatments.

## Acknowledgements

This work was supported by the Intramural Research Program of the National Institute of Neurological Disorders and Stroke (NINDS), NIH. We acknowledge the Children’s Brain Tumor Network (CBTN) for providing access to critical research data. We thank Jane Jones and the NIH Protein Synthesis Core for their contributions to the synthesis and optimization of Angubindin-1. We also extend our gratitude to Eric Bunker for his assistance with AlphaFold analyses.

