## Supplemental Figures for "Angulin-1/LSR inhibition transiently disrupts the blood-tumor barrier to enhance doxil permeability and impair malignant glioma progression"

**Supplementary Materials and Methods**

*Mass Spectrometry (LC/MS)*

Mass spectrometry data were acquired on an Agilent 6100 Series Quadrupole LC/MS System, (Agilent Technologies, Inc., Santa Clara, CA, USA) equipped with electrospray source, operated in the positive-ion mode. Separation was performed on 300SB-C3 Poroshell column (Agilent Technologies, 2.1 mm x 75 mm; particle size 5 □m). Mass spectra were recorded across the range 300–3000 m/z. The UV signal was collected at 280 nm with a reference at 360 nm. Data acquisition and analysis were performed using OpenLAB CDS ChemStation Edition C.01.05.

*Circular Dichroism*

Circular dichroism measurements for protein secondary structure determination were performed on J-1500 CD spectrophotometer (Jasco Inc., Easton, MD, USA). Samples (concentration 1 mg/ml in HEPES buffer) were run in the interval 240 – 190 nm at 25oC in continuous mode at scanning speed 50 nm/min, digital integration time 8 seconds, bandwidth 1 nm. Due to low signal/noise ratio in the spectral area below 200 nm, cylindrical demountable quartz cuvettes of 20 mm diameter and pathlength 0.5, 0.2, 01 and 0.01 mm (Hellma USA, New York, USA) were used. The spectra of the buffer were collected under identical conditions and were subtracted from the sample spectra. The evaluation of protein secondary structure was performed using online Dichroweb server supported by Birkbeck College, University of London, UK. The fit was performed using Selcon 3 (Self-Consistent Method) algorithm and the Reference Set 7 (48 proteins with known secondary structure). The results were represented as percentage content of helical, beta-sheet, and unoriented elements.

*Dynamic Light Scattering*

Protein's particle sizes and distributions were assessed by dynamic light scattering using a DynaPro Plate Reader (Wyatt Technology, Santa Barbara, CA, USA). Before measurement the sample was spun with tabletop centrifuge for 5 minutes at 13,200 rpm. Twenty microliters of protein solution (1 mg/mL) in HEPES + NaCl + TCEP buffer was added to a 384-well plate, illuminated with an 830 nm laser, and light scattering was measured at 22 °C. Each measurement consisted of 20 acquisitions, with each acquisition lasting 5 seconds. To obtain the hydrodynamic radii the intensity autocorrelation functions were fitted by a regularization algorithm using Dynamics software version 8.10 (Wyatt Technology, Santa Barbara, CA, USA).

*Differential Scanning Fluorimetry*

The thermal unfolding and refolding of a protein was studied with differential scanning fluorometer Prometheus NT.48 (NanoTemper Technologies GmbH, Munich, Germany). The excitation wavelength was 285 nm, the scanning speed was 1 degree/min., the temperature interval was 20 – 90 degrees Celsius. The collected signals were fluorescence emission at 330 and 350 nm, their ratio and back-scattering.

*Mass Photometry*

The mass photometry experiments were performed on mass photometer TwoMP (Refeyn Ltd., Oxford, UK).  The data acquisition was performed with AcquireMP (version 2023 R1.1) software and data analysis was performed with DiscoverMP (version 2023 R1.2) software.

The instrument calibration was performed with a mix of beta-amylase from sweet potato (Sigma-Aldrich, St. Louis, MO, USA) and thyroglobulin from bovine thyroid (Sigma-Aldrich, St. Louis, MO, USA) dissolved in HEPES buffer. The droplet-dilution procedure of probe preparation was applied. The autofocusing was performed with pure buffer added to the gasket well, then the drop of protein solution was added and thoroughly mixed by pipette pumping to achieve final well content concentration of 24 to 100 nM. The recording time was 60 seconds. The experiments were triplicated with preparation of new probes each time. The data analysis was performed in histogram mode with manual selection of observed peaks and application of Gaussian fit.

**<u>Supplementary Figure Legends</u>**

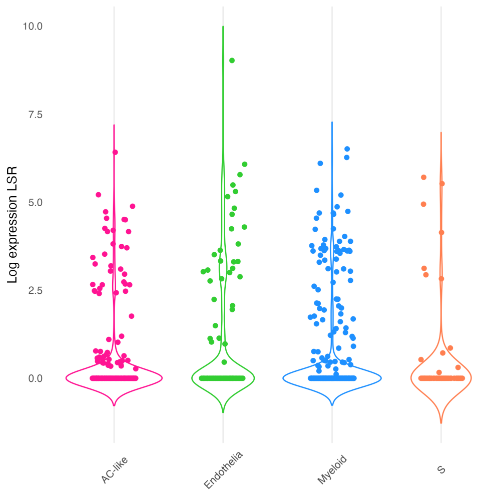


<u>Supplementary Figure 1:</u> Violin plot of LSR expression by cell type focused on cell types with expression greater than zero noted in dot plot (Figure YC) (AC-like (pink), Endothelia (green), Myeloid (blue), S (orange)). Expression levels were log-transformed using the log1p function to accommodate zeros and normalize the distribution.

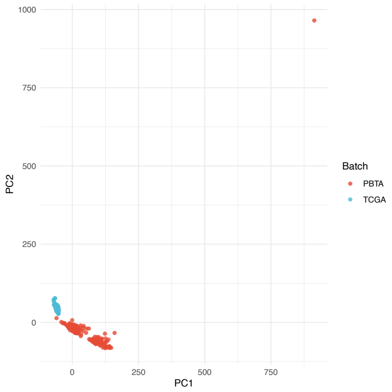


Supplementary Figure 2: PCA plot showing the separation of PBTA and TCGA samples before batch correction. Each point represents a sample, colored by batch TCGA or PBTA. One sample from PBTA in the top right corner was noted as an outlier (C4115334). (PBTA = red, TCGA = blue); TCGA (n=157) and PBTA (n=258).

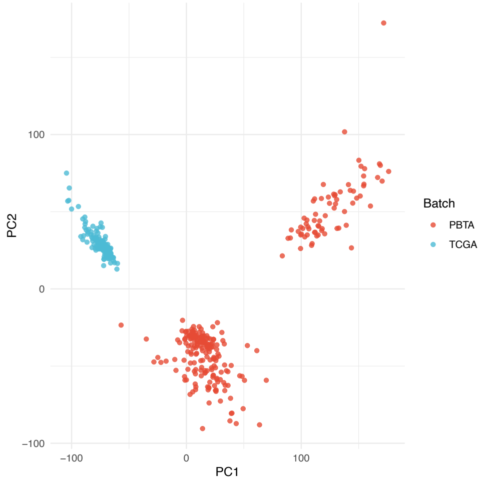


Supplementary Figure 3: PCA plot showing the separation of PBTA and TCGA samples after removing outlier (C4115334). Each point represents a sample, colored by batch; (PBTA = red, TCGA = blue); TCGA (n=157) and PBTA (n=257).

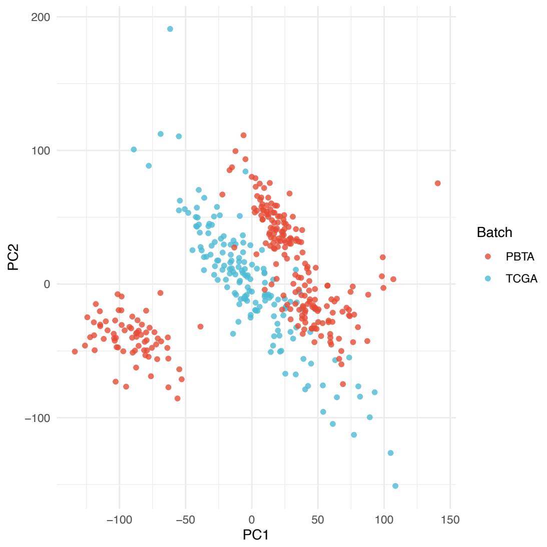


Supplementary Figure 4: PCA plot showing the separation of PBTA and TCGA samples after batch correction. Each point represents a sample, colored by batch (TCGA or PBTA); (PBTA = red, TCGA = blue); TCGA (n=156) and PBTA (n=257). One outlier from PBTA (C4115334) and one patient from TCGA without recorded age (TCGA-28-2510) were removed to facilitate analyses involving patient age.

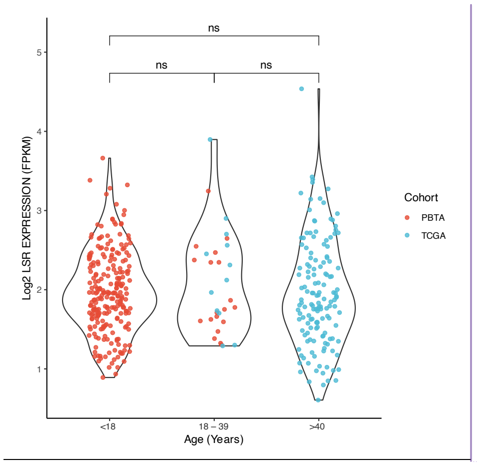


Supplementary Figure 5: Violin plot comparing LSR expression (log-transformed) across different age groups (under 18y, 18 – 39y, 40y and over) in TCGA (n=156) and PBTA (n=257)

**Commented [MOU1]:** Can you check with Yang to ensure the X axis is correct? “18…x<40) and …40 seems weird cohorts, one outlier from PBTA (C4115334) and one patient from TCGA without recorded age (TCGA-28-2510) were removed from the analysis. Points are jittered and colored by cohort (PBTA = red, TCGA = blue); ns: not significant with p>0.05.

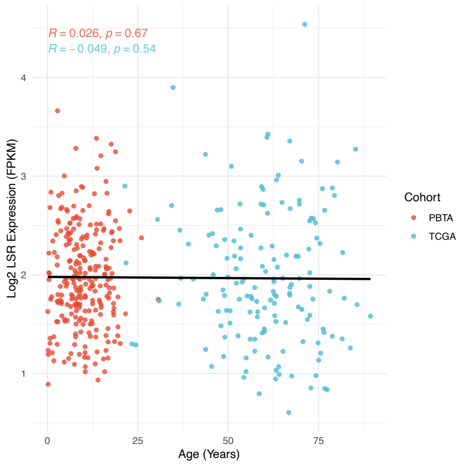


Supplementary Figure 6: Scatter plot showing the correlation between LSR expression and age (in years) across all samples. The plot includes a regression line and Pearson correlation coefficient. The dots are colored by cohort (PBTA = red, TCGA = blue). Pearson correlation coefficient R and p value is listed at the left top panel and samples are colored by cohort, TCGA (n=156) and PBTA (n=257), cohorts, one outlier from PBTA (C4115334) and one patient from TCGA without recorded age (TCGA-28-2510) were removed from the analysis.

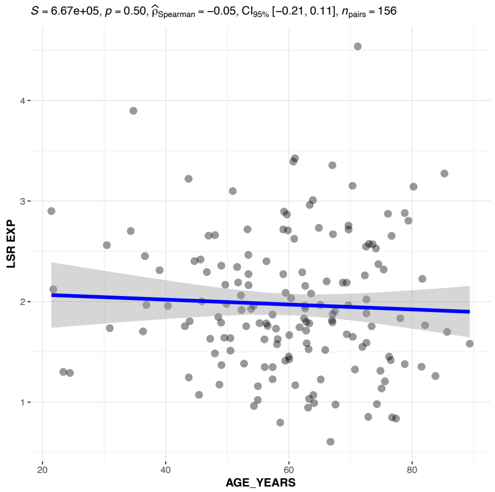


Supplementary Figure 7: Scatter plot showing the correlation between LSR expression and age in TCGA samples TCGA (n=156). One patient from TCGA without recorded age (TCGA-28-2510) was removed from the analysis. The plot includes a regression line and Pearson correlation coefficient.

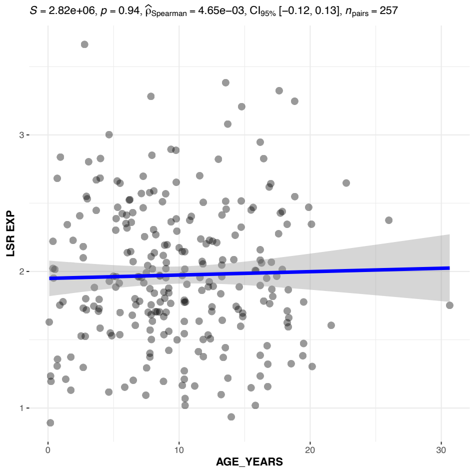


Supplementary Figure 8: Scatter plot showing the correlation between LSR expression and age in PBTA samples. One outlier from PBTA (C4115334) was removed from the analysis. The plot includes a regression line and Pearson correlation coefficient. PBTA (n=257).

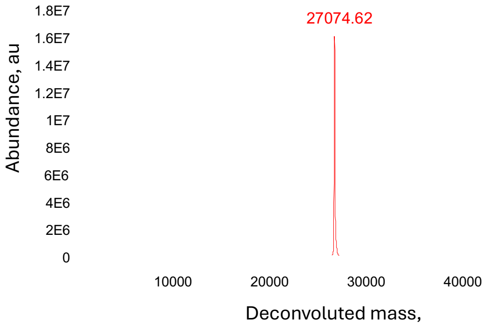


Supplementary Figure 9: Deconvoluted mass spectrum of Angubindin-1. Chromatogram revealed a single, dominant peak with a relative abundance of 100%; corresponding to a molecular weight of 27,074.62 Da, confirming high purity and expected mass of the protein.

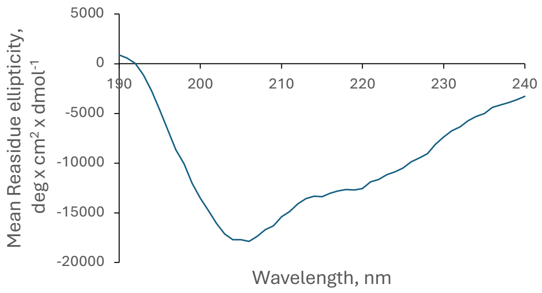


Supplementary Figure 10: The circular dichroism (CD) spectrum of Angubindin-1 solution in HEPES buffer, optical path 0.01 mm. CD spectra displayed characteristic features of a mixed α-helical and β-sheet conformation, including a pronounced negative peak in the 203–208 nm range indicative of α-helical elements. Angubindin-1 adopts a well-ordered conformation in solution under the tested conditions.

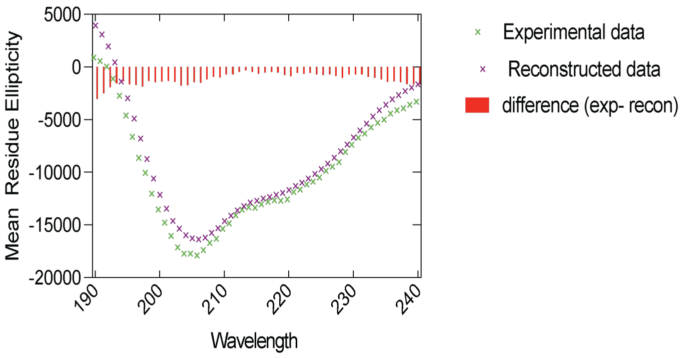


Supplementary Figure 11: The comparative CD-spectra, raw data (green patterns) and DichroWeb-reconstructed data (purple patterns); the parameters of fitting are presented in Materials and Methods session. The red bars represent the residuals (difference between raw and reconstructed data).

To assess the purity and molecular integrity of Angubindin-1, we analyzed it using liquid chromatography–mass spectrometry (LC-MS). The resulting chromatogram revealed a single, dominant peak with a relative abundance of 100%; corresponding to a molecular weight of 27,074.62 Da, confirming high purity and expected mass of the protein (Supplemental Figure 9). The secondary structure of the purified recombinant Angubindin-1 was evaluated using circular dichroism (CD) spectroscopy. The CD spectra displayed characteristic features of a mixed α-helical and β-sheet conformation, including a pronounced negative peak in the 203–208 nm range indicative of α-helical content (Supplemental Figures 10-11). Spectral data were deconvoluted using the DichroWeb online server [1], yielding the following estimates of secondary structure composition:

| Structure Type | Percentage |
|---|---|
| α-helices | 29.2% |
| β-sheets | 31.1% |
| β-turns | 28.6% |
| Disordered | 12.9% |

Supplementary Table 1: Spectral deconvolution data showing the percentage of α-helices, β-sheets, β-turns, and disordered regions in Angubindin-1, derived from CD analysis using DichroWeb. Minor deviation from 100% total is attributed to limitations in spectral fitting, as commonly observed in CD deconvolution analyses.

These results suggest that Angubindin-1 adopts a well-ordered conformation in solution under the tested conditions. Minor deviation from 100% total is attributed to limitations in spectral fitting, as commonly observed in CD deconvolution analyses [1]. To evaluate the thermal stability of Angubindin-1's secondary structure, differential scanning fluorimetry (Nano-DSF) was performed. This method monitors intrinsic tryptophan and tyrosine fluorescence changes in response to temperature [2]. As shown in Supplemental Figure 12, the thermal ramping revealed a clear transition with an unfolding (melting) temperature of 65.6 °C (±0.18) and a closely matching refolding temperature of 65.3 °C (±0.05), suggesting reversible folding and a stable secondary structure supported by balanced enthalpic and entropic contributions. Given that protein multimerization can influence biological function, the oligomeric status of Angubindin-1 was assessed using dynamic light scattering (DLS). At 1 mg/mL (37 µM) in HEPES buffer, DLS revealed a highly homogeneous population (polydispersity index: 11.5%) with a hydrodynamic radius of 3.4 nm, consistent with a molecular weight of ~58.2 kDa, indicating a predominantly dimeric state under these conditions (Supplemental Figure 13).

To further assess concentration-dependent oligomerization, mass photometry was used to analyze Angubindin-1 at submicromolar concentrations [3]. At 24 nM, the protein appeared predominantly monomeric, with a mass distribution centered at ~33 kDa (±8.4 kDa). As concentrations increased (30–100 nM), the mass distribution shifted upward and broadened, consistent with dimer formation. At 100 nM, a peak at ~58 kDa confirmed a predominance of dimeric species suggesting that Angubindin-1 exists in a dynamic monomer-dimer equilibrium depending on concentration (Supplemental Figure 14). To determine the molecular weight and oligomerization state of Angubindin-1, we performed mass photometry across a range of concentrations. At low concentrations (~20 nM), Angubindin-1 appeared predominantly monomeric, as indicated by a sharp, well-defined peak corresponding to its monomeric mass. However, as the concentration increased (30–100 nM), we observed a notable broadening and rightward shift of the mass distribution, suggesting a concentration-dependent transition toward dimer formation. These results indicate that Angubindin-1 exists primarily as a monomer at low concentrations but forms dimers or higher-order multimers within the working concentration range.  At the 600 µg/mL dose employed in this study, Angubindin-1 is predominantly dimeric, highlighting a dynamic oligomerization profile that could shape its biological activity.

**Commented [MOU2]:** Based on the concentration difference, is 600ug/mL a high or low dose of angubindin-1? Would be good to have a good tie in sentence of whether we're treating our cells and animals with high or low dose angubindin-1 (monomeric vs. dimeric conformation of this agent).

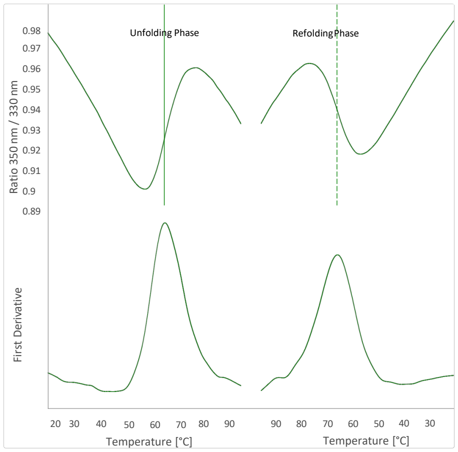


Supplementary Figure 12: The experimental nano-DSF thermograms of unfolding and refolding of Angubindin-1. The upper plots represent the ratio of fluorescence emission intensities at 350

nm and 330 nm; the bottom ones represent the first derivative of these signals. The thermal ramping revealed a clear transition with an unfolding (melting) temperature of 65.6 °C (±0.18) and a closely matching refolding temperature of 65.3 °C (±0.05), suggesting reversible folding and a stable secondary structure supported by balanced enthalpic and entropic contributions

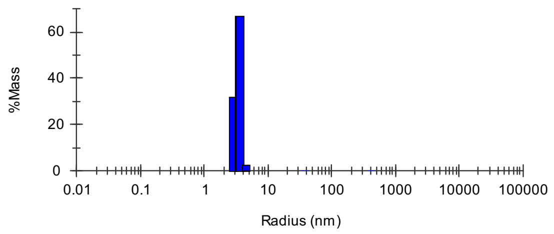


Supplementary Figure 13: The size-distribution of Angubindin-1 in 1 mg/ml HEPES solution by dynamic light scattering with a peak at 3.4 nm, corresponding to 58.2 kDa molecular weight. At 1 mg/mL (37 µM) in HEPES buffer, DLS revealed a highly homogeneous population (polydispersity index: 11.5%) with a hydrodynamic radius of 3.4 nm, consistent with a molecular weight of ~58.2 kDa, indicating a predominantly dimeric state under these conditions.

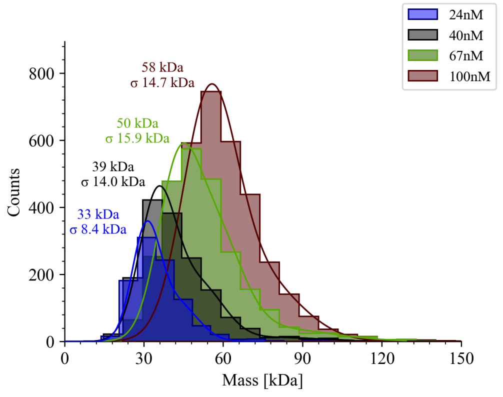


Supplementary Figure 14: Mass-photometry analysis of association status of Angubindin-1 at variable concentrations in HEPES buffer. The peak positions indicate molecular weight and corresponding standard deviation. At 24 nM, the protein appeared predominantly monomeric, with a mass distribution centered at ~33 kDa (±8.4 kDa). As concentrations increased the mass distribution shifted upward and broadened, consistent with dimer formation. At 100 nM, a peak at ~58 kDa confirmed a predominance of dimeric species suggesting that Angubindin-1 exists in a dynamic monomer-dimer equilibrium depending on concentration. At low concentrations (~20 nM), Angubindin-1 appeared predominantly monomeric, as indicated by a sharp, well-defined peak corresponding to its monomeric mass. However, as the concentration increased (30–100 nM), we observed a notable broadening and rightward shift of the mass distribution, suggesting a concentration-dependent transition toward dimer formation.

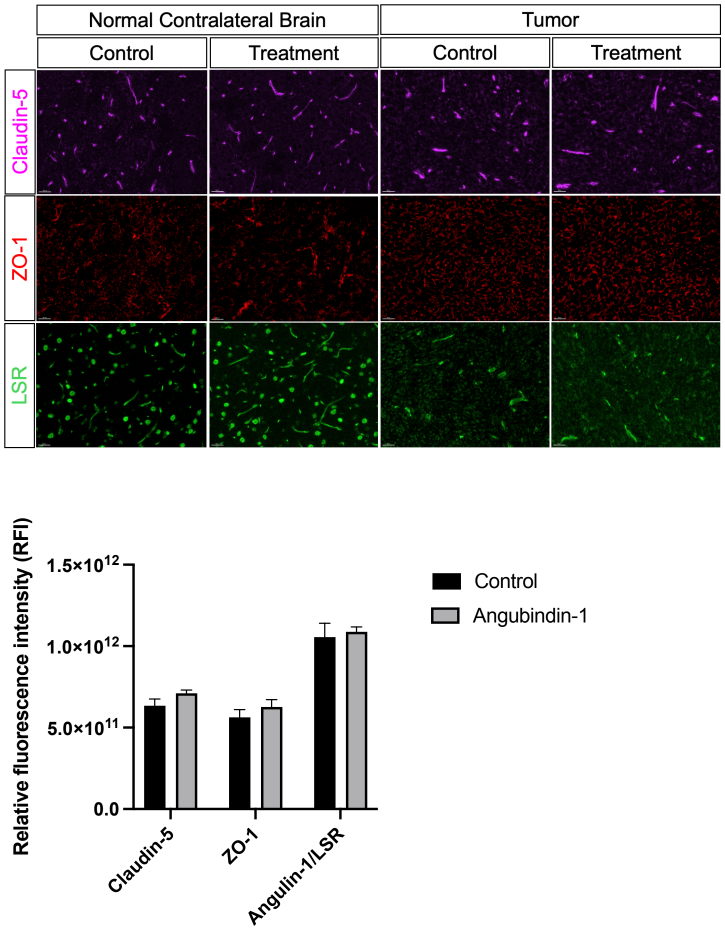


Supplementary Figure 15: No difference in tight junction expression post Angubindin-1 measured after two doses on day 14. Immunofluorescent qualitative staining for claudin-5, ZO-1 and LSR within normal contralateral brain vs. tumor tissue and quantitative relative fluorescence of the entire brain slice.
