## Supplemental Table 1 for "Angulin-1/LSR inhibition transiently disrupts the blood-tumor barrier to enhance doxil permeability and impair malignant glioma progression"

| Structure Type | Percentage |
|---|---|
| α-helices | 29.2% |
| β-sheets | 31.1% |
| β-turns | 28.6% |
| Disordered | 12.9% |

**Supplementary Table 1. Estimated Secondary Structure Composition of Angubindin-1 Based on Circular Dichroism Spectroscopy.** Spectral deconvolution data showing the percentage of α-helices, β-sheets, β-turns, and disordered regions in Angubindin-1, derived from CD analysis using DichroWeb.
